# On the cortical mapping function – visual space, cortical space, and crowding

**DOI:** 10.1101/621458

**Authors:** Hans Strasburger

## Abstract

The retino-cortical visual pathway is retinotopically organized: Neighbourhood relationships on the retina are preserved in the mapping to the cortex. Size relationships in that mapping are also highly regular: The size of a patch in the visual field that maps onto a cortical patch of fixed size follows, along any radius and in a wide range, simply a linear function with retinal eccentricity. As a consequence, and under simplifying assumptions, the mapping of retinal to cortical locations follows a logarithmic function along that radius. While this has already been shown by Fischer (1973), the link between the linear function – which describes the local behaviour by the cortical magnification factor *M* – and the logarithmic location function for the global behaviour, has never been made fully explicit. The present paper provides such a link as a set of ready-to-use equations using Levi and Klein’s E_2_ nomenclature, and examples for their validity and applicability in the retinotopic mapping literature are discussed. The equations allow estimating *M* in the retinotopic centre and values thus derived from the literature are provided. A new structural parameter, *d*_*2*_, is proposed to characterize the cortical map, as a cortical counterpart to *E*_*2*_, and typical values for it are given. One pitfall is discussed and spelt out as a set of equations, namely the common myth that a pure logarithmic function will give an adequate map: The popular omission of a constant term renders the equations ill defined in, and around, the retinotopic centre. The correct equations are finally extended to describe the cortical map of Bouma’s law on visual crowding. The result contradicts recent suggestions that critical crowding distance corresponds to a constant cortical distance.

## Introduction

One of the most beautiful organizational principles of the human brain is that of topographical mapping. Whilst perhaps universal to the brain, its regularity is most apparent for the three primary senses mediated through the thalamus – sight, hearing, and touch – i.e., in retinotopy, tonotopy, and somatotopy. For the visual domain with which we are concerned here, the regularity of topography is particularly striking and is at a level that lends itself to mathematical description by analytic functions. The seminal papers by Fischer (1973) and Schwartz (1977, 1980) derive the complex logarithm as a suitable function for mapping the location in the visual field to the location of its projection’s in (a flat-map of) the primary visual cortex, by which the visual field’s polar-coordinate grid gets mapped onto a rectilinear cortical grid. The log function’s image domain – the complex plane – is reinterpreted thereby as a two-dimensional real plane.^1^ As Schwartz explains in the two papers, the rationale for employing the log function in the radial direction is that its first derivative is an inverse-linear function, the latter implicit in the cortical magnification concept for the visual field as proposed by Daniel & Whitteridge (1961). Expressed more directly, the *integral* of an inverse linear function is the logarithmic function. Intuitively, summing-up (integrating over) little steps on the cortical map, where each step obeys cortical magnification, will result in the log mapping.

Schwartz’s (1977, 1980) papers with the complex-log mapping have become rather popular in visual psychophysics and visual neurophysiology. Van Essen, Newsome & Maunsell (1984), e.g., use it for explaining the topography of the macaque’s primary visual cortex, writing “Along the axis corresponding to constant polar angle, magnification is inversely proportional to eccentricity, and hence distance is proportional to the logarithm of eccentricity (x ∝ log E)” (p. 437). Levi, Klein & Aitsebaomo (1985, Fig. 14) and Virsu et al. (1987, Fig. 7) plot psychophysical thresholds in terms of cortical units. As another example, Klein & Levi (1987), in the context of modelling hyperacuity in peripheral vision, derive from the log rule that, if vernier-acuity offsets are assumed to have a constant cortical representation – i.e. one that is independent of eccentricity – vernier offsets will depend linearly on eccentricity in the visual field (we will come back to that in the last section). Horton & Hoyt (1991) use it to point out that the well-known inverse-linear function for the cortical magnification factor *M* (CMF) follows from a log-spaced cortical map. Engel et al. (1997, Fig. 9, Fig 12; 1994, Fig. 2), and Larsson & Heeger (2006), use the (real-valued) log function implicitly when they use an exponential for the inverse location function (which corresponds to a log forward mapping). Duncan & Boynton (2003) fit their fMRI activity maps for the V1 topology using Schwartz’s complex-log mapping. The most advanced development is Schira, Tyler, Spehar & Breakspear’s (2010) closed-form analytic representation for the cortical maps, at the same time accommodating for the horizontal-vertical anisotropy and preserving cortical area constancy across meridians by an added shear function.

While Fischer’s and Schwartz’s papers present the mathematical relationships (withexamples for their application) Klein & Levi (1987) provide an *empirical* link between psychophysical data and location on the cortical map. For characterizing the inverse-linear CMF-vs-eccentricity function, they use a concept they had developed earlier for psychophysical results (Levi, Klein, & Aitsebaomo, 1984; Levi et al., 1985): The slope of that linear function, when normalized to the foveal value, can be quantified by a single number, called *E*_*2*_. The concept is illustrated graphically in Figure 1B below: In an x-y plot vs eccentricity, *E*_*2*_ is the (negative) X-axis intercept or, alternatively, the (positive) eccentricity value at which the foveal value is incremented by itself (i.e., doubles). Klein & Levi (1987) further bridge the gap to proportionality when they show that relationships become simpler and more accurate when the data are not treated as a function of eccentricity *E* itself, but of a transformed eccentricity, *E\**, referred to as *effective eccentricity, E* = E + E*_*2*_. The *linear* cortical magnification function thereby turns into *proportionality*. In the cortical map, locations – i.e. distances from the retinotopic centre – are then proportional to the logarithm of effective eccentricity, x ∝ log *(E+E*_*2*_). The approach is verified by showing the empirical data both as thresholds and in cortical units (Klein & Levi, 1987, Fig. 5; for rescaling that figure’s right ordinate the authors posit that 1 mm of cortex corresponds to ~10% of effective eccentricity).

**Figure 1.**
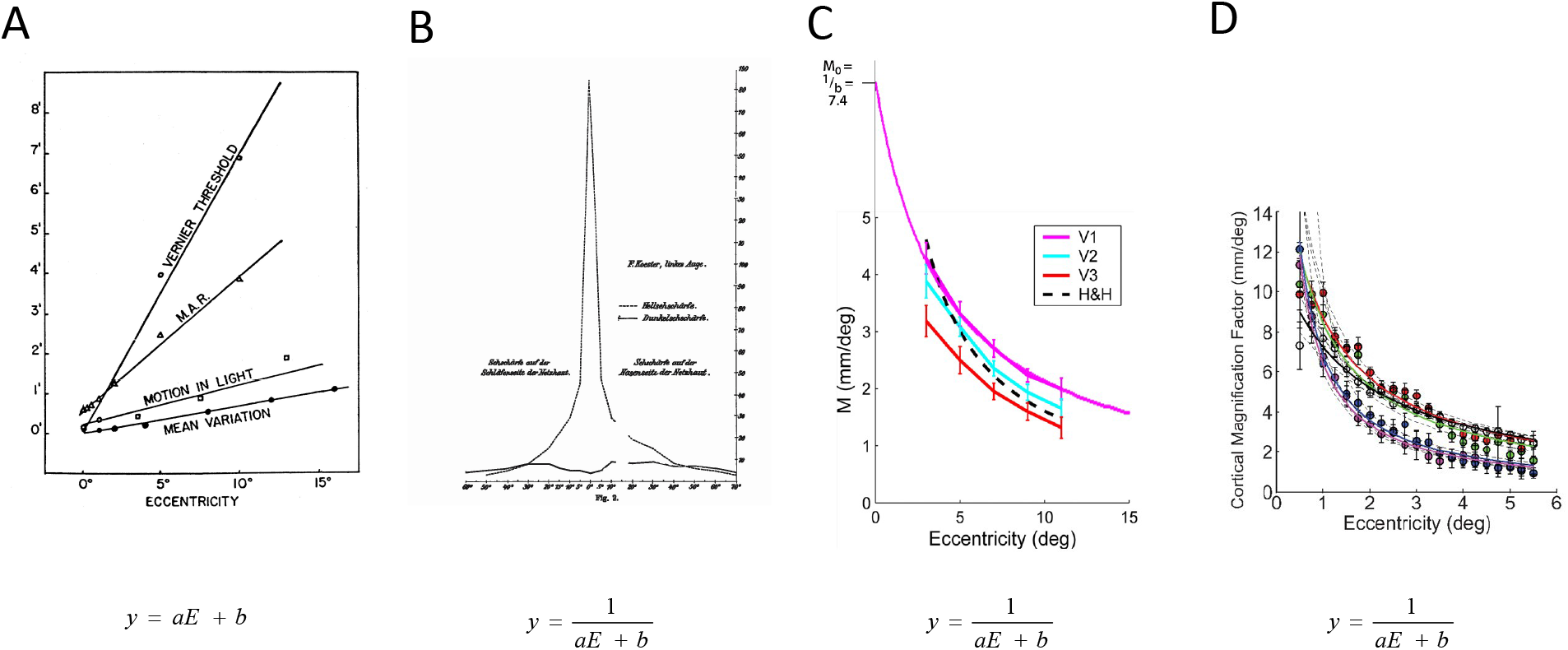
Examples for the linear and the inverse-linear (approx. hyperbola) graph. Even though the two are equivalent, their intuitive interpretation is often different, with the linear graph taken as evidence of a shallow performance decline and the inverse-linear graph as evidence of a steep decline (Rosenholtz, 2016; Strasburger 2020). A. MAR for various visual performance parameters; Weymouth (1958, Fig. 13). B. Visual acuity; Fick (1898, Fig. 2). C. Cortical magnification factor *M*; Dougherty et al. (2003, Fig. 5). A hyperbola graph, obtained from linear regression to the inverse data (*M*=1/(0.033*E*+0.1355)), and an axis intercept *M*_*0*_=1/*b* have been added to the original graph. D. Same; Harvey & Dumoulin, 2011, Fig. 4B. In C and D note the steep incline toward the retinotopic centre and that no data are obtained in or near the centre. The central value *M*_*0*_ is therefore difficult to derive directly from those graphs.

However, the papers discussed so far have not yet fully exploited the tight mathematical link between the linear and the logarithmic law for its empirical use. While the basic mathematical form of the mapping function − log *(E) or* log *(E+E*_*2*_) − is drawn upon and made use of, further parameters are left free to vary and to be determined by fitting to the data. The derivations in the present paper take the log-mapping approach one step further. Unlike these and other papers (discussed below), the parameters for the logarithmic map are here obtained by mathematical derivation from the linear law. In a neuroscience context, that law will be the inverse of the CMF. For the psychophysicist, measures of low-level visual-perceptual function like the minimal angle of resolution (MAR) can be an approximation. In both cases, Levi and Klein’s *E*_*2*_ concept is the basis here. We thereby arrive at a set of fully explicit equations that allow converting the linear, local-behaviour law of the CMF, specified by *E*_*2*_, to a description of the global behaviour, the *location* on the cortical map. These equations are the message of the paper. In a next step, the empirical data for the cortical maps (from fMRI or single-cell analysis) are then used to verify the correctness of those parametrical equations. This approach represents a more principled one than before. It further places additional constraints on the describing functions, thus adding to their reliability.

Since such derivations have been attempted before and have led to erroneous results or have stopped short of exploring the implications, derivations are presented in a step-by-step manner, considering at each step what that means. Key equations are highlighted by surrounding boxes for easy spotting, i.e. those that should be of practical use in describing the cortical map. Or, for example, for obtaining improved estimates for the foveal CMF, *M*_*0*_.

Instead of the complex log we here consider the simpler case of the real-valued, 1D mapping, where *eccentricity* in the visual field, expressed in degrees of visual angle along a radius, is mapped onto the *distance* of its representation from the retinotopic centre, expressed in millimetres. The resulting real-valued logarithmic function shall be called the *cortical location function*. Taking the 1D case implies no loss of generality; the function is easily generalized to the 2D case by writing it as a vector function. Compared to the complex log, the real function has the added advantage of allowing separate parameters for the horizontal and vertical meridian, required to meet the visual field’s horizontal-vertical anisotropy.

Once these relationships for the cortical location function are established, they need to be verified by empirical data. We use data from the literature and our own for this. It turns out that not only do the fits work excellently, and even better than the original fits, but that the constraints imposed by the parametric equations can also be used for the long-standing problem of improving estimates for the foveal CMF (*M*_*0*_). Another issue addressed there are attempts to become independent of the retinotopic centre’s location. That centre’s exact location appears to be difficult to find (it is often specified only approximately), and some authors like to use some other reference location instead. It turns out, however, that while equations can be referenced to some other location than the centre, true independence from the latter cannot be achieved by any means.

In the context of these derivations, I propose a new metric, *d*_*2*_, measured in millimetres, for characterizing the cortical map. It is the equivalent of *E*_*2*_ (which is measured in degrees visual angle). Like *E*_*2*_ in the visual field, *d*_*2*_ allows specifying the steepness of location change in the cortical map, e.g. for quantifying the horizontal-vertical anisotropy or even for comparisons between species.

In another section, it is further argued that the simplified version (x ∝ log *E*) that is not uncommon in the fMRI literature needs to be avoided and that the full version with a constant term added in *the log’s argument* needs to be employed (i.e., x ∝ log (*E* + c)). There is further apparently confusion about what does and what does not represent the required constant term, which adds to a common myth that omitting the term simplifies matters.

Finally, the cortical location function can be used, perhaps unexpectedly, to derive the cortical distances in visual crowding. Crowding happens when neighbouring patterns to a target stimulus are closer than a critical distance; that critical distance can be described by Bouma’s law (Bouma, 1970; Strasburger, Harvey, & Rentschler, 1991; Pelli, Palomares, & Majaj, 2004; Pelli & Tillman, 2008; Whitney & Levi, 2011; Strasburger, 2020). We thus arrive at a cortical version of Bouma’s law. While this has been done before (Levi et al., 1985; Motter & Simoni, 2007; Pelli, 2008; Nandy & Tjan, 2012; Strasburger, Rentschler, & Jüttner, 2011; Strasburger & Malania, 2013), the present derivations go beyond those in that they include the fovea and provide the derivations.

## 1. Concepts

Peripheral vision is unlike central vision as Ptolemy (90–168) already noted. Yet just how it is different is still a puzzling question. The goal here is to draw the attention to the highly systematic organization of the early neural processing stages by deriving equations that describe its architecture. But before doing so we need to be explicit on a number of concepts that are the foundation for what follows.

### The linear law and the hyperbola graph

Four types of analytic functions are central for describing functional dependencies on eccentricity – in the visual field or in retinotopic areas: linear and inverse-linear, and logarithmic and exponential. Their graphs look entirely different (giving rise to misleading intuition; Rosenholtz, 2016, Strasburger, 2020) yet the first two and second two are effectively equivalent to each other. Let’s start with the first pair (the second pair follows in Figure 3).

Aubert and Foerster’s (1857) characterization of the performance decline with retinal eccentricity as a linear increase of minimum resolvable size – sometimes referred to as the Aubert-Foerster law – is still the conceptual standard. It corresponds to what is now called *M*-*scaling* when based on cortical magnification (Virsu & Rovamo, 1979; Virsu et al., 1987) or the change of *local spatial scale* when the scaling factor is not thus constrained (Watson, 1987). Examples for the linear law are shown in Figure 1A and 2A. However, by the end of the 19^th^ century it also became popular to use the inverse of minimum size instead, i.e. acuity, in an attempt to make the sensory decline more graphic (e.g. Fick, 1898, shown in Figure 1B). And, since the inverse-linear function’s graph is close to a hyperbola, we arrive at the well-known hyperbola-like function of, e.g., acuity vs. eccentricity seen in most textbooks, or in Østerberg’s (1935) equally well-known cone-density graph. Examples of that graph for the cortical map are in Dougherty et al. (2003, Fig. 5) and Harvey & Dumoulin (2011, Fig. 4B), shown in Figure 1 C and D.

Yet, graphic as it may be, the hyperbola graph does not lend itself to a comparison of decline parameters. Weymouth (1958) therefore already argued for using the linear graph, introducing the concept of the *minimal angle of resolution* (MAR) as a general measure of size threshold. Weymouth summarized how the MAR and other spatial visual performance parameters depend on retinal eccentricity (Figure 1A). Importantly, Weymouth stressed the *mandatory use* of a non-zero, *positive y-axis intercept* for these functions (Weymouth, 1958, p. 109). This will be a major point here in the paper; it is related to the necessity of a constant term in the cortical-location function as discussed below.

### Cortical magnification

Daniel & Whitteridge (1961) and Cowey & Rolls (1974) introduced cortical magnification as a quantitative concept for retinotopic mapping, which, for a given visual-field location, summarizes functional density along the retino-cortical pathway into a single number. The linear cortical magnification factor (CMF), *M,* was defined as the *diameter in the primary visual cortex onto which 1 deg of the visual field projects* (*areal M* was defined as an areal counterpart). Enlarging peripherally presented stimuli by *M* turns out to counter performance decline to a large degree for many visual tasks (reviewed, e.g., by Virsu et al., 1987); it was thus suggested as a general means of equalizing visual performance across the visual field (Rovamo & Virsu, 1979). Even though this so-called strong hypothesis was soon dismissed (e.g. Westheimer, 1982, p. 161), the strong tie between cortical distances and (in particular) low-level psychophysical tasks is still striking.

The relationship between the early visual architecture and psychophysical tasks is still a matter of debate; why, for example, do different visual tasks show widely differing slopes of their eccentricity functions (Figure 1A)? In contrast, the manner in which the CMF varies with eccentricity is largely agreed upon: *M* decreases with eccentricity – following approximately a hyperbola (Figure 1C and D) – and its inverse increases linearly (Schwartz, 1980; Van Essen et al., 1984; Tolhurst & Ling, 1988; Horton & Hoyt, 1991, Slotnick, Klein, Carney, & Sutter, 2001, Duncan & Boynton, 2003; Larsson & Heeger, 2006; Schira, Wade, & Tyler, 2007). Figure 2A shows a few examples for the latter. Note that in the figure there is one function from psychophysics shown along with the anatomical estimates (Rovamo & Virsu, 1979; Virsu & Rovamo, 1979; Virsu et al., 1987). Note also that all functions need to have a positive y-axis intercept, *be it ever so slight,* because otherwise *M* were undefined, i.e., infinite.

**Figure 2.**
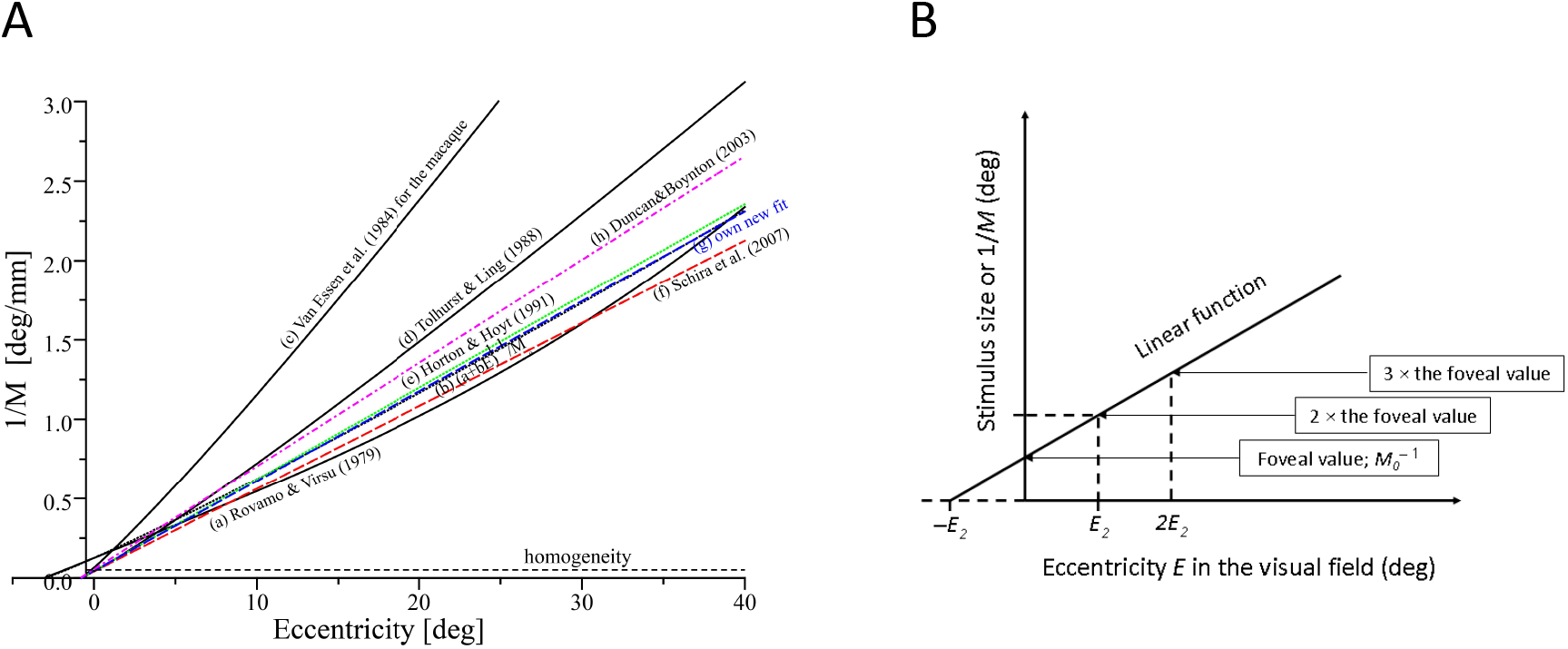
A. The inverse of the cortical magnification factor or, equivalently, the size of a patch in the visual field that projects onto a patch of constant size in the cortex, as a function of eccentricity in the visual field (Fig. 9 in Strasburger et al., 2011, reproduced for illustrating the text). All functions show a mostly linear behaviour. Their slope is quite similar, with the exception of Van Essen et al.’s (1984) data for the macaque; other data show similar slopes between human and monkey (e.g. Oehler, 1985). Note that Rovamo & Virsu’s function is based on psychophysical data. Note also that all functions need to have a positive y-axis intercept. B. An illustration of the *E*_*2*_ concept. *E*_*2*_ is defined as the eccentricity where the foveal value doubles, or (equivalently) as the eccentricity increment that leads to an increment by the foveal value. It is also the negative x-axis intercept. Note that the foveal value does not double *every E*_*2*_ increment (cf. Strasburger, 2020). Importntly, note that the concept can be used for both psychophysical and anatomical data.

### Other equations

Empirical data typically fit the linear concept quite well in the considered range of about 40° eccentricity, but, nevertheless, fits can sometimes be improved by introducing a slight nonlinearity (Table 1). Rovamo, Virsu, & Näsänen (1978), as an example, used a polynomial by adding a small 3^rd^-order term; Van Essen et al. (1984), Tolhurst & Ling (1988), and Sereno et al. (1995) increased the exponent of the linear term slightly above 1. Virsu & Hari (1996) used a sine function, based on geometrical considerations. Only a part of the sine’s period comes into play so that the function is still close to linear in that range. The latter function is interesting because it is the only one that can be extended to eccentricities larger than 90°(cf. Strasburger, 2020). However, improvements over a linear approach are mostly small or absent and do not warrant the added complexity in the derivations to follow, so we will not pursue this further.

**Table 1.**
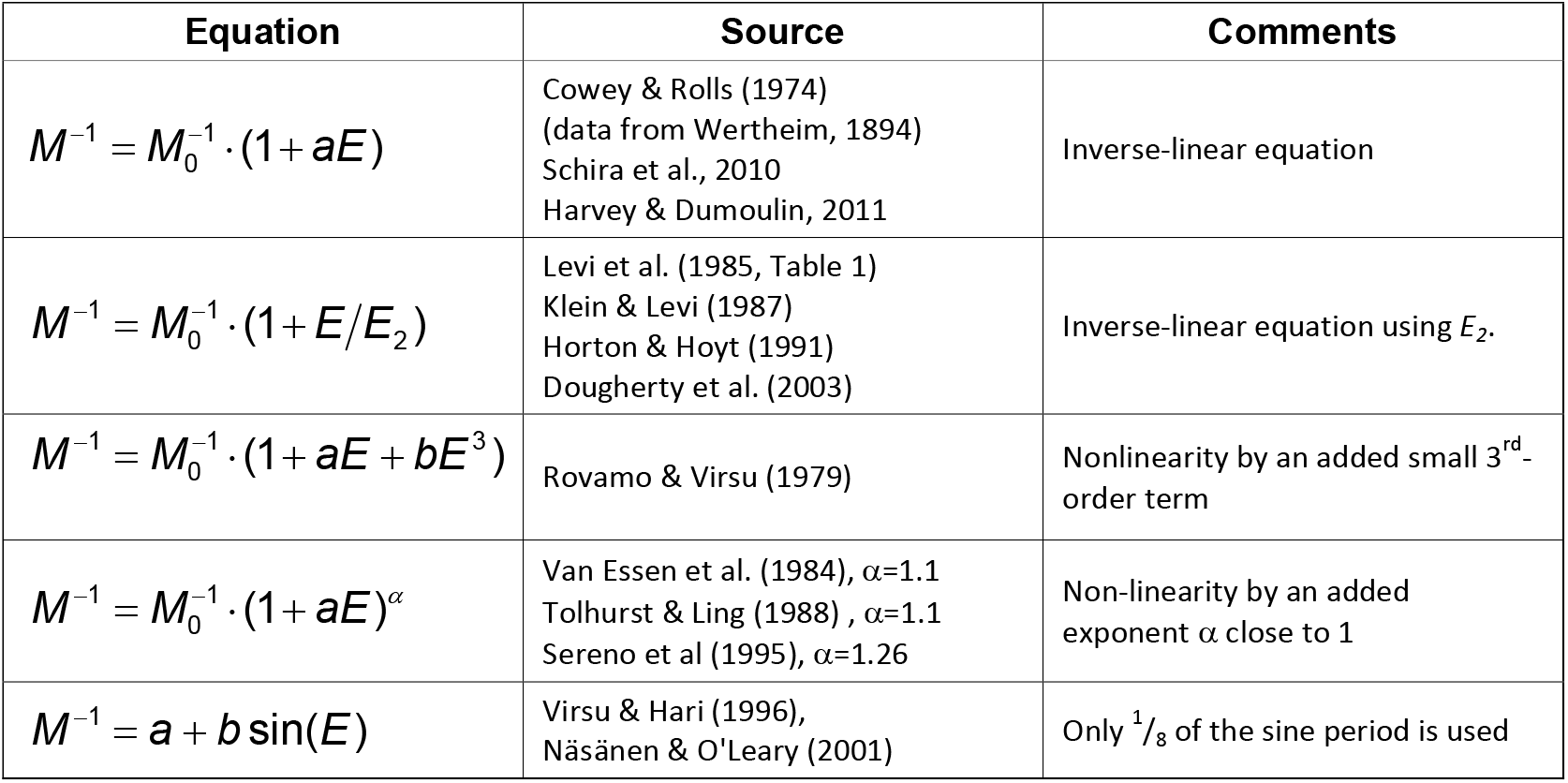
Equations used for describing eccentricity functions (modified from Strasburger et al., 2011).

### The E_2_ concept

For a quick comparison of eccentricity functions, Levi et al. (1984, p. 794) introduced the *E*_*2*_ concept by pointing out the specific eccentricity at which the respective foveal value doubles (Figure 2B). More generally, *E*_*2*_ is the *eccentricity increment* at which y increases by the foveal value. I.e., at eccentricity *E*_*2*_ the foveal value is doubled and at twice *E*_*2*_ is tripled. As a graphic aide, *E*_*2*_ is also the distance from the origin of where the linear function crosses the eccentricity axis.

*E*_*2*_ is most often used for psychophysical tasks but lends itself equally well for describing the anatomical function (Levi et al., 1985, Table 1; Klein & Levi, 1987; Horton & Hoyt, 1991; Dougherty et al., 2003). Eq. (1) states the corresponding equation.

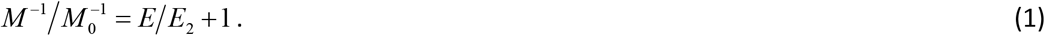

*M*^−1^ in that equation is measured in °/mm (one might call it the retinal magnification factor: it corresponds to the receptive field size of a cortical neuron on the retina). 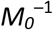 is that value in the fovea’s centre. The function’s slope is given by 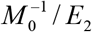, so when these functions are normalized to the foveal value, their slope is 1/*E*_*2*_. I.e., larger *E*_*2*_ corresponds to shallower slope. Parameter *E*_*2*_ thus captures an important property of the functions (how they increase/decrease) in a single number. A summary of values was reported by Levi et al. (1984), Levi et al. (1985), Klein & Levi (1987), or more recently by Strasburger et al. (2011, Tables 4–6). These reported *E*_*2*_ values vary widely between different visual functions. They also vary considerably for functions that seem directly comparable to each other (for example, *E*_*2*_ for vernier acuity: 0.62°–0.8°; for *M*^*−1*^: 0.77°–0.82° or even 3.67° in Dougherty et al., 2003; for Landolt-C acuity: 1.0°–2.6°; letter acuity: 2.3°–3.3°; gratings: 2.5°–3.0°). On the other hand, *E*_*2*_ can also be surprisingly similar for tasks that seem entirely unrelated, like for example the *E*_*2*_ of 1.22° for the perceived travel extent in the fine-grain movement illusion (Foster, Thorson, McIlwain, & Biederman-Thorson, 1981). Note also the limitations of *E*_*2*_: since, for example, the empirical functions always deviate a little from linearity, the characterization by *E*_*2*_, by its definition, works best at small eccentricities.

### M-scaling and local scale

The left hand ratio in eq. (1), 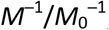, is the ratio by which a peripherally seen stimulus needs to be size-scaled to occupy cortical space equal to a foveal stimulus. So the equation can be re-written as

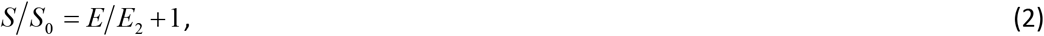

where *S* is *scaled size* and *S*_*0*_ is the size at the fovea’s centre. *S*_*0*_ can be considered the size-scaling unit in the visual field, and *E*_*2*_ the locational scaling unit (i.e. the unit in which scaled eccentricities are measured). If *E*_*2*_ refers to the cortical map, this is the *concept of M-scaling*. If *E*_*2*_ in the equation refers to some other eccentricity function, this corresponds to a more abstract way of size scaling, called *local scale* (Watson, 1987).

### The cortical location function

Fischer (1973) and Schwartz (1977, 1980) proposed the complex log function for mapping the visual field to the cortical area. The key property of interest for that mapping is the behaviour *along a radius* (from the fovea) in the visual field; the simpler real-valued log function can thus be used instead of the complex logarithm. This, then, maps the eccentricity in the visual field to the distance from the retinotopic centre on the cortical map (Figure 3B). Neuroscience papers often prefer to show the inverse function (i.e. mirrored along the diagonal with the x and y axis interchanged, thus going “backwards” from cortical distance to eccentricity), which is the exponential function shown schematically in Figure 3A.

**Figure 3.**
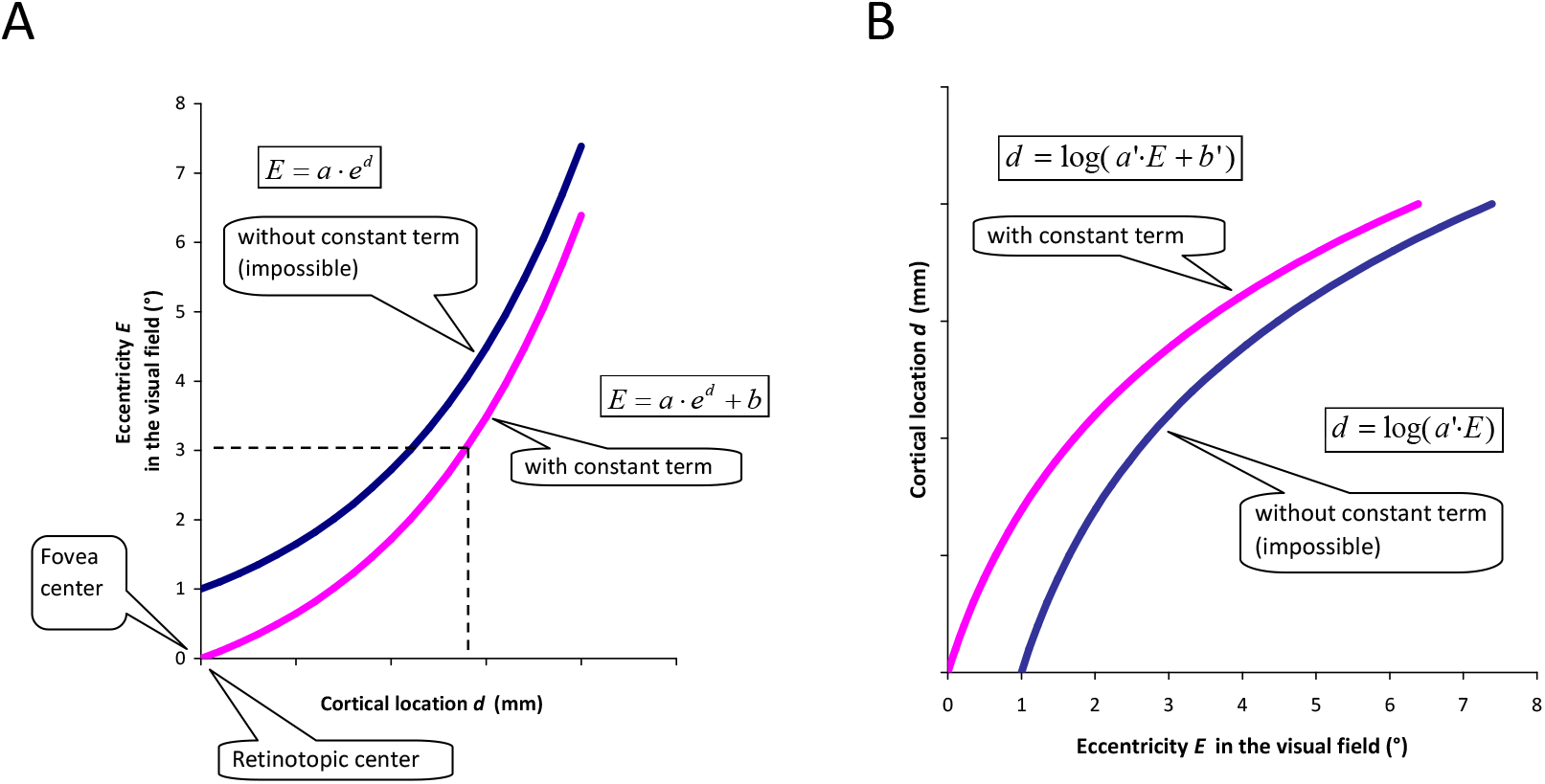
Schematic graph of the cortical location function introduced by Fischer (1973) and Schwartz (1977, 1980), along a radius from the retinotopic centre. A version with, and another without a constant term (parameter *b* or *b’* in the equation) are shown. The constant term’s omission was intended as a simplification for large eccentricities but is not physically possible near or in the foveal centre. The graph in shows eccentricity *E* as a function of cortical distance *d* (which is an exponential); Schwartz (1980) discussed mainly the inverse function shown in (B), i.e. for *d* as a function of *E* (which is logarithmic).

### The constant term

Schwartz (1980) has discussed two versions of the function that differ in whether there is a constant term added in the argument; the difference is illustrated in the graph. The version without the constant is often considered simpler and is thus often (inappropriately) preferred. A point in the following will be that that simplicity is deceiving and can lead to wrong conclusions – and more complicated equations. Note that the constant term is at different places in the equations: For the exponential function in figure part (A) it is *added to* the exponential, for the logarithmic function in (B) it is *within* the log’s argument. As will be seen later, the constant term in both cases corresponds to the positive y-intercept of the linear function (Figure 2B).

### The retinal and the retinotopic centre

There is an important difference in difficulty between measuring at the fovea’s exact centre and at the cortical retinotopic counterpart. Whereas psychophysical measurements at the fovea are particularly simple and reliable, determining the exact retinotopic centre and the CMF at that location, *M*_*0*_, appear the most difficult and *M*_*0*_’s value is mostly extrapolated from peripheral values. The consequences of this include different strategies in research between the two fields regarding the map.

### Anisotropy

The visual field is not isotropic: Performance declines differently between radii. Slopes differ between vertical and horizontal, and upper vs lower field. Accordingly, iso-performance lines (for the binocular field) are distorted ellipses rather than circular outside the central visual field, which is isotropic (e.g. Wertheim, 1894, Harvey & Pöppel, 1972; Pöppel & Harvey, 1973). Rovamo & Virsu (1979, p. 498) accordingly computed separate *M* estimates for each meridian. There is further a nonlinearity at the transition from the isotropic to the anisotropic field (Pöppel & Harvey, 1973, Fig. 6). Correspondingly, early visual areas are also anisotropic (e.g. Horton & Hoyt, 1991). The effect of anisotropy on the cortical magnification factor is quantitatively treated by Schira et al. (2007, 2010); their *M*_*0*_ estimate is the geometric mean of the isopolar and isoeccentric *M* estimates. In the equations presented below, the horizontal/vertical anisotropy can be accommodated by letting the parameters depend on the radius in question. There are further anisotropies that are not accounted for by varying slopes along the radii (Schira et al., 2007, 2010). These authors, for preserving *area* constancy across meridians, thus extend modelling by a shear function (using the hyperbolic secans; Schira et al., 2010, eq. 6 and Fig. 2). Mappings then differ between meridians, with deviations from linearity most noticeable on, and close to, the vertical meridian at around 1° eccentricity (Schira et al., 2010, Fig. 2). The derivations presented below, for simplicity, do not include these refinements.

### Symbols in the paper

To keep the overview, symbols used in the paper are summarized in Table 2. Some of those are in standard use and some are newly introduced in the remainder.

**Table 2.**
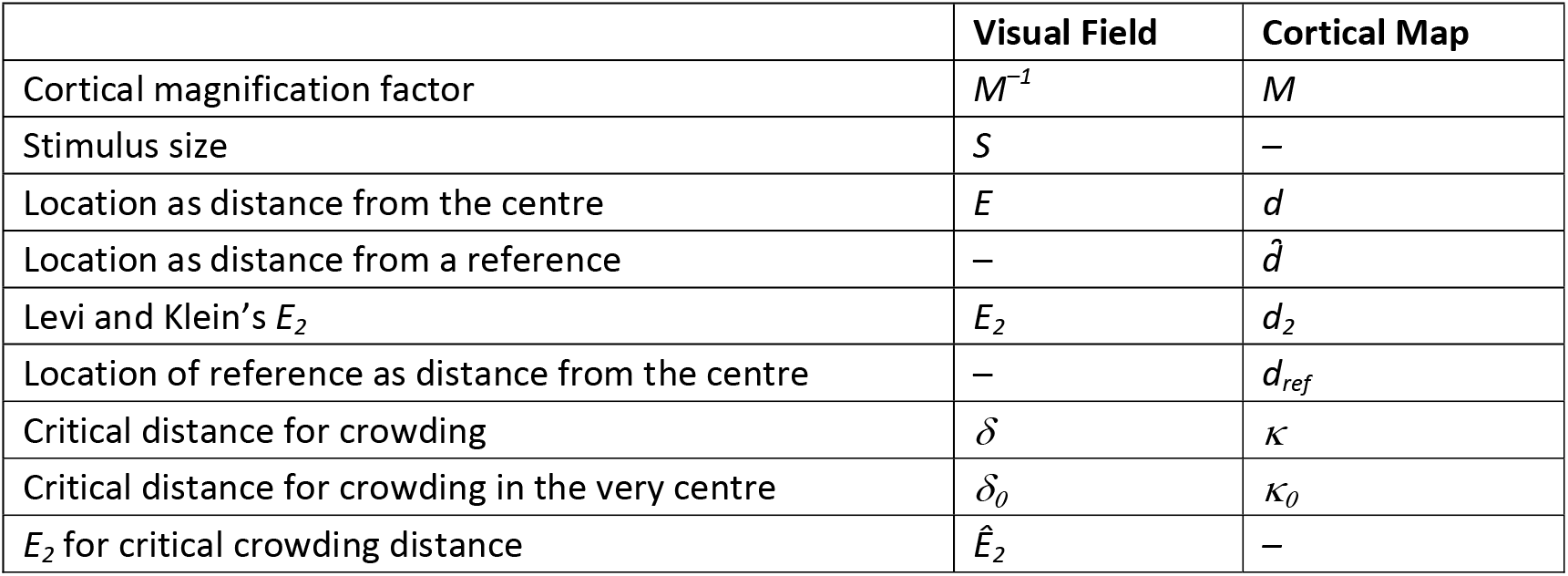
Summary of symbols used in the paper

|  | Visual Field | Cortical Map |
| --- | --- | --- |
| Cortical magnification factor | $M^{-1}$ | $M$ |
| Stimulus size | $S$ | – |
| Location as distance from the centre | $E$ | $d$ |
| Location as distance from a reference | – | $\hat{d}$ |
| Levi and Klein's $E_2$ | $E_2$ | $d_2$ |
| Location of reference as distance from the centre | – | $d_{ref}$ |
| Critical distance for crowding | $\delta$ | $\kappa$ |
| Critical distance for crowding in the very centre | $\delta_0$ | $\kappa_0$ |
| $E_2$ for critical crowding distance | $\hat{E}_2$ | – |

## 2. The cortical location function

### 2.1 Cortical location specified relative to the retinotopic centre

The ratio *S/S*_*0*_ in eq. (2) is readily estimated in psychophysical experiments as the size of a stimulus relative to its foveal value for achieving equal perceptual performance. However, its physiological counterpart 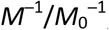 in eq. (1) appears difficult to assess directly, even though it is a physiological concept. Instead, it is typically derived by extrapolation from peripheral values, e.g. from the cortical-location function *d* = *d(E)* (Figure 3). The function links a cortical distance *d* in a retinotopic area to the corresponding distance in the visual field that it represents. More specifically, *d* is the distance (in mm) on the cortical surface between the representation of a visual-field point at eccentricity *E*, and the representation of the fovea centre. Under the assumption of linearity of the cortical magnification function *M*^*−1*^*(E)*, this function is logarithmic (Figure 3B) and its inverse *E = E(d)* exponential (Figure 3A), as shown by Fischer (1973) and Schwartz (1977, 1980). Since the *E*_*2*_ parameter allows a simple formulation of the linear eccentricity functions (Figure 2), as e.g. in eq. (1), it will be useful to state the location function with those notations. First steps have been derived in Strasburger et. al. (2011, eqs. 10 – 13; corresponding here eqs. 3 – 6). The present derivations go further. The location function allows a concise quantitative characterization of the early retinotopic maps.

For its derivation, notice first that, locally, the cortical distance of the respective representations *d(E)* and *d(E+ΔE)* of two nearby points along a radius, at eccentricities *E* and *E+ΔE,* is given by *M(E)·ΔE.* This follows from *M*’s definition and the fact that *M* refers to 1°. The cortical magnification factor *M* is thus the first derivative of *d(E)*, i.e.,

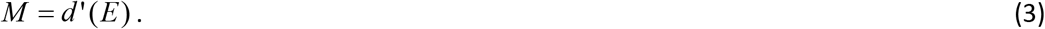

Conversely, the location *d* on the cortical surface (i.e., the global aspect) is the integral over *M,* starting at the fovea centre:

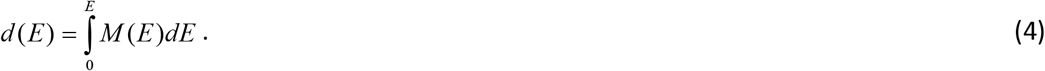

If we insert eq. (1) – i.e. the equation using *E*_*2*_ – into eq. (4), we have

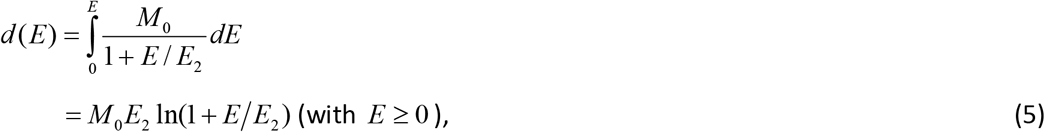

where *ln* denotes the natural logarithm.

The inverse function, *E(d)* is derived by inverting eq. (5),

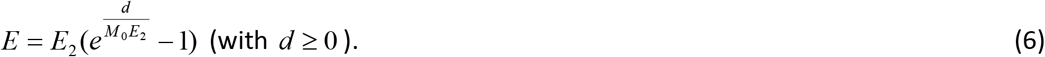

It states how the eccentricity *E* in the visual field depends on the distance *d* of the corresponding location in a retinotopic area from the retinotopic centre. With slight variations, discussed below, it is the formulation often referenced in fMRI papers on the cortical mapping. Note that, by its nature, it is only meaningful for positive values of cortical distance *d*. The significance of this point will become apparent later.

We can simplify that function further by introducing an analogue to *E*_*2*_ in the cortex. Observe that like any point in the visual field the location at *E*_*2*_ has a representation (on the meridian in question), whose distance from the retinotopic centre we denote as *d*_*2*_. Thus, *d*_*2*_ in the cortex represents *E*_*2*_ in the visual field.

To express eq. (6) using *d*_*2*_ instead of *M*_*0*_, first apply the equation to that location *d*_*2*_:

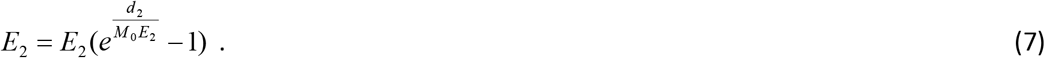

Solving that for the product *M*_*0*_ *E*_*2*_ gives

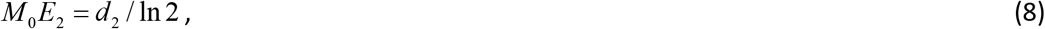

which, inserted into eq. (6) in turn gives

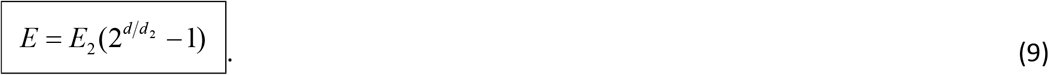

Eq. (9) is the most concise way of stating the cortical location function. We can also restate it however as

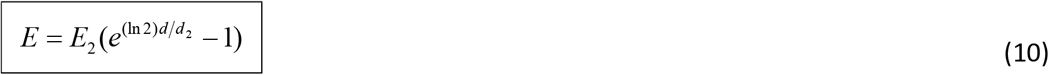

since the exponential to the base e is often more convenient (*ln* again denotes the natural logarithm).

This equation (eq. 10) is particularly nice and simple provided that *d*_*2*_, the cortical equivalent of *E*_*2*_, is known. That value, *d*_*2*_, could thus play a key role in characterizing the cortical map, similar to the role of *E*_*2*_ in visual psychophysics (cf. Table 4 – Table 6 in Strasburger et al., 2011, or earlier the tables in Levi et al., 1984, Levi et al., 1985, or Klein & Levi, 1987). Estimates for *d*_*2*_ derived from literature data are summarized in Section 2.4 below, as an aid for concisely formulating the cortical location function.

The new cortical parameter *d*_*2*_ can be calculated from eq. (8), restated here for convenience:

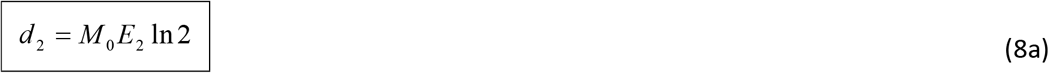

### 2.2 Cortical location specified relative to a reference location

Implicit in the definition of *d* or *d*_*2*_ is the knowledge about the location of the fovea centre’s cortical representation, i.e. of the retinotopic centre. That locus has proven to be hard to determine precisely, However, and instead of the centre it has thus become customary to use some fixed eccentricity *E*_*ref*_ as a reference. Engel et al. (1997, Fig. 9; 1994, Fig. 2), for example, use *E*_*ref*_ = 10°. Larsson & Heeger (2006, Fig. 5) use *E*_*ref*_ = 3°.

To restate eq. (6) or (10) accordingly, i.e. with some reference eccentricity different from *E*_*ref*_ = 0, we first apply eq. (10) to that reference:

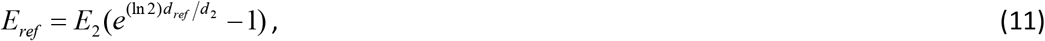

where *d*_*ref*_ denotes the value of *d* at the chosen reference eccentricity, e.g. at 3° or 10°.

Solving then that equation for *d*_*2*_ and plugging the result into eq. (9) or (10), we arrive at

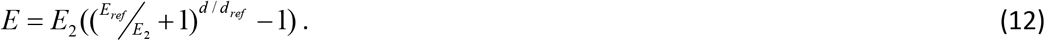

Expressed to the base e instead, we have

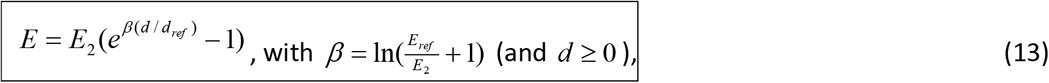

which represents the location function expressed *relative to a reference eccentricity E*_*ref*_, and its equivalent in the cortical map, *d*_*ref*_. (One could also derive eq. (13) directly from eq. (6).) Note that if, in that equation, *E*_*2*_ is taken as the reference eccentricity for checking, it reduces to eq. (10) as expected. So, *E*_*2*_ can be considered as a special case of a reference eccentricity. Note further that, unlike the location equations often used in the retinotopy literature (Van Essen et al., 1984, in the introduction; Duncan & Boynton, 2003; Larsson & Heeger, 2006), the equations are well defined in the fovea centre: for *d* = 0, the eccentricity *E* is zero, as it should.

What reference to choose is up to the experimenter. However, the fovea centre itself cannot be used as a reference eccentricity – the equation is undefined for *d*_*ref*_ = 0 (since the exponent is then infinite). Thus, the desired independence of knowing the retinotopic centre’s location has not been achieved − that knowledge is still needed, since *d,* and *d*_*ref*_, in these equations are defined as the respective distances from that point.

Equations (12) and (13) have the ratio *d/d*_*ref*_ in the exponent. It is a proportionality factor for cortical distance. From the intercept theorem in geometry we know that this factor cannot be re-expressed by any other expression that leaves the zero point undefined. True independence from knowing the retinotopic centre, though desirable, thus cannot be achieved.

We can nevertheless shift the coordinate system such that locations are specified relative to the reference location, *d*_*ref*_. For this, we define a new variable 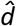 as the cortical distance (in mm) from the reference *d*_*ref*_ instead of from the retinotopic centre (see Figure 4 for an illustration for the shift and the involved parameters), where *d*_*ref*_ is the location corresponding to some eccentricity, *E*_*ref*_. By definition, then,

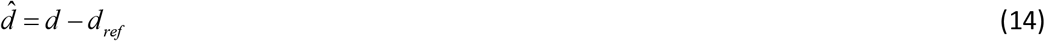

**Figure 4:**
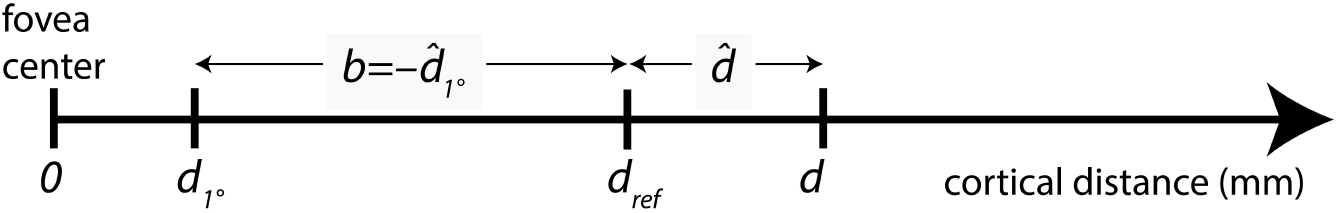
Illustration of the cortical distance measures used in equations (6) – (23), and of parameter *b* in eq. (18) further below. *d* – cortical distance of some location from the retinotopic centre, in mm; *d*_*ref*_ – distance (from the centre) of the reference that corresponds to *E*_*ref*_; *d*_*1°*_ – distance of the location that corresponds to *E = 1°*; 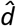 – distance of location *d* from the reference *d*_*ref*_.

In the shifted system – i.e., with 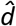 instead of *d* as the independent variable – eq. (6) for example becomes

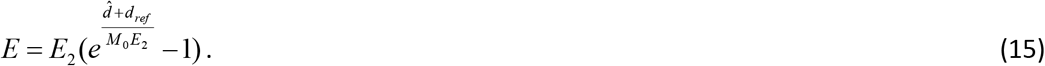

The equation might be of limited practical use, however (like eq. 6 from which it was derived), since the parameters *M*_*0*_ and *E*_*2*_ in it are not independent; they are inversely related to each other as seen in eq. (8) or (8a) (or eq. 17). That interdependency is removed in eq. (9) or (10), (which work from the retinotopic centre), or eq. (13) (which used a reference eccentricity). The latter (eq. 13), in the shifted system becomes

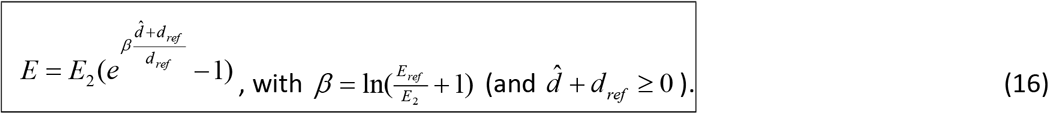

That equation now has the advantage over eq. (15) of having only two free parameters, *E*_*2*_ and *d*_*ref*_. (*E*_*ref*_ is not truly free since it is empirically linked to *d*_*ref*_.) The foveal magnification factor *M*_*0*_ has dropped from the equation. Indeed, by comparing eq. (13) to eq. (6) (or by comparing eq. (15) to (16)), *M*_*0*_ can be calculated from *d*_*ref*_ and *E*_*2*_ as

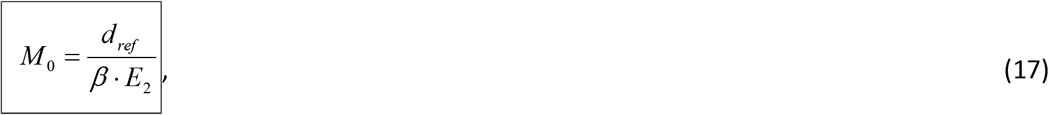

where *β* is defined as in the previous equation. With an approximate location of the retinotopic centre (needed for calculating *d*_*ref*_) and an estimate of *E*_*2*_, that latter equation leads to an estimate of the foveal magnification factor, *M*_*0*_ (see Section 2.4 for examples).

Equations (16) and (17) are crucial to determining the retinotopic map in early areas. They should work well for areas V1 to V4 as discussed below. The connection between the linear and log or exponential function based estimations provided by these equations allows cross-validating the empirically found parameters and thus leads to more reliable results. Duncan & Boynton (2003), for example, review the linear law and also determine the cortical location function empirically but do not draw the connection. Their’s and others’ approaches are discussed as practical examples in the section after next (Section 2.4).

### 2.3 Independence from the retinotopic centre with the simplified function?

Schwartz (1980) had offered a simplified location function where the constant term is omitted, which works at sufficiently large eccentricities. Frequently that was the preferred one by other authors as seemingly being more practical. The present section briefly highlights how this approach leads astray if pursued rigorously.

The simplified version of the location function 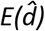 omits the constant term in eq. (6) and those that follow from it (i.e., the “−1” in eq. 6 up to eq. 16). Instead, the equation

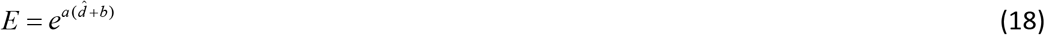

is fit to the empirical data, with free parameters *a* and *b*. The distance variable in it is 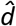 as before, i.e., the cortical distance from a reference *d*_*ref*_ representing some eccentricity *E*_*ref*_ in the visual field. Engel et al. (1997, Fig. 9; 1994, Fig. 2), for example, use *E*_*ref*_ = 10° for such a reference, and for that condition report the equation 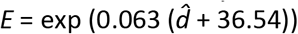. Larsson & Heeger (2006, Fig. 5), as another example, use *E*_*ref*_ = 3°, and for area V1 in that figure give the function 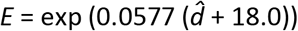. Note that neither of these equations contains the required constant term (cf. Figure 3), since the constants (36.54 and 18.0) are inside, not outside the exponential’s argument.

**Figure 5.**
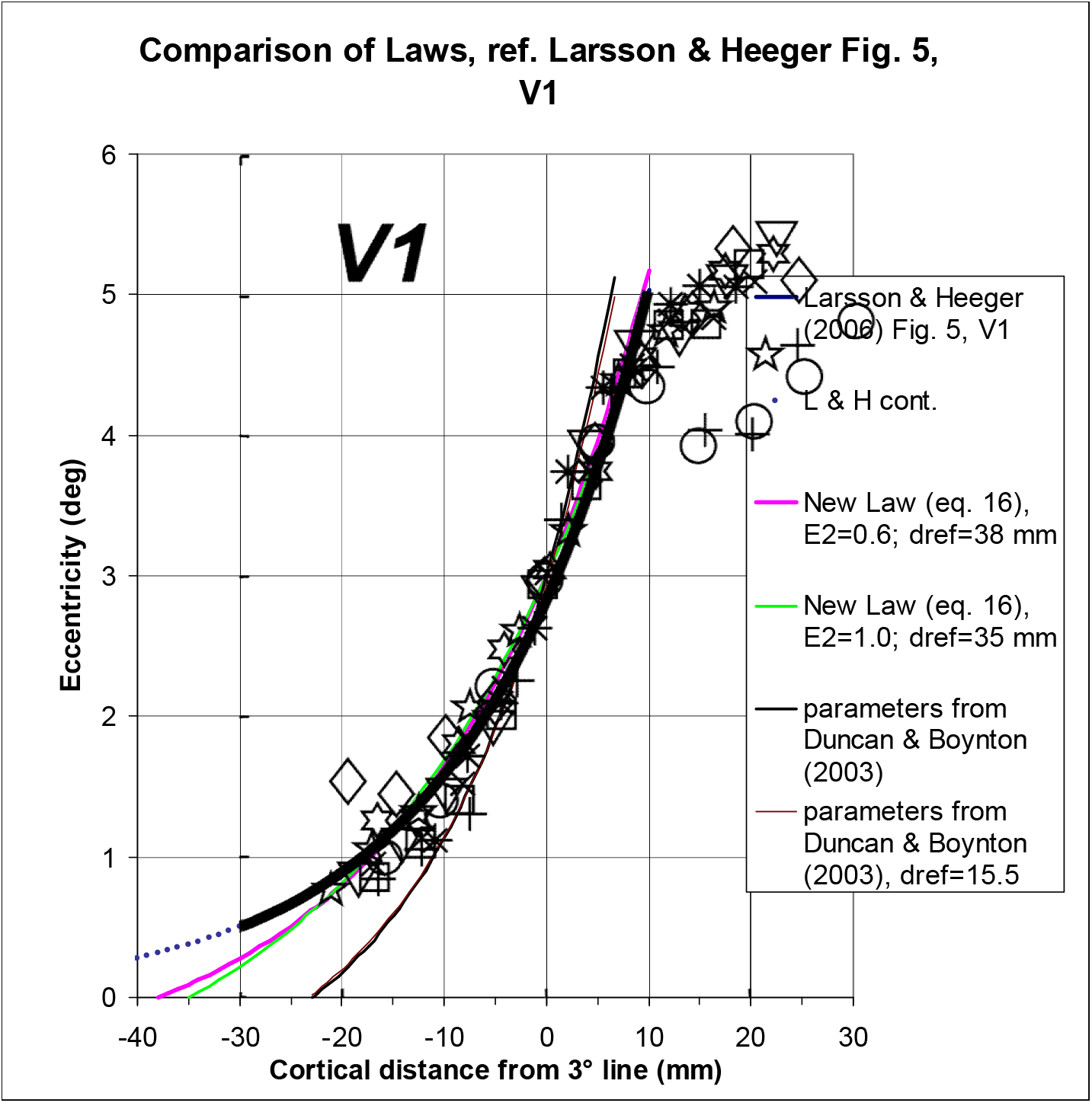
Comparison of conventional and improved analytic functions for describing the cortical location function. Symbols show the retinotopic data for area V1 with reference location *d*_*ref*_ = 3° from Larsson and Heeger (2006, Fig. 5) (symbols for nine subjects). Superimposed is the original fit (thick black line), according to eq. (18) 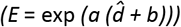 or eq. (19), i.e. a fit without a constant term). The blue dotted line continues that fit to lower eccentricities; the fitted 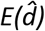 function goes to (negative) infinite cortical distance, which is physically meaningless. The pink and green line show graphs of the preferable eq. (16) that was derived from integrating the inverse linear law (eq. 1).The equations are underconstrained if *M*_*0*_ is not known; two pairs of parameter choices are shown, [*E*_*2*_ = 0.6°, *d*_*ref*_ = 38 mm] and [*E*_*2*_ = 1.0°, *d*_*ref*_ = 35 mm], respectively. The corresponding retinotopic centre’s magnification factor *M*_*0*_ can be calculated by eq. (17) as 35.4 mm/° and 25.3 mm/° for the two cases, respectively. Black and brown line: 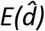 function with parameters derived by Duncan & Boynton (2003), *M*_*0*_ = 18.5 mm/° and *E*_*2*_ = 0.831° (black), and with *d*_*ref*_ = 15.5 mm (brown) for comparison (discussed in the next section). Note that, by definition, the curves from Larsson & Heeger pass through the 3° point at 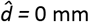. Note also that, according to the authors, the data beyond ~10 mm were biased and can be disregarded.

We can attach meaning to the parameters *a* and *b* in eq. (18) by constraining the function appropriately (see Strasburger, 2019, for the derivation). By that we arrive at an equation

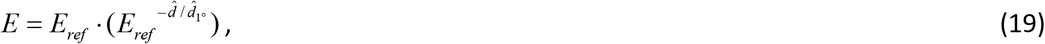

where 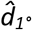 is the distance of the 1° line from the reference eccentricity’s representation; it is around −36.5 mm for *E*_*ref*_ = 10° as used by Engel et al. (1994, 1997).

This is now the *simplified* cortical location function, i.e. the simplified analogue to eq. (16), with parameters spelt out. One can easily verify that the equation holds true at the two defining points, i.e. at 1° and the reference eccentricity. Note also that, as intended, knowing the retinotopic centre’s location in the cortex is not required since 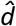 is defined relative to a non-zero reference. However, in between these two points the function has the wrong curvature (see Fig. 4 in the next section, fat black line). Importantly, however, the equation fails with small eccentricities, for the simple reason that *E* cannot become zero in that equation. In other words, the fovea’s centre is never reached, even at the retinotopic centre.

So the seeming simplicity of eq. (18) that we started out from leads astray in and around the fovea – which, after all, is of prime importance for vision. The next section illustrates the impact of the constant term with data from the literature.

### 2.4 Practical use of the equations: examples

#### 2.4.1 The approach of Larsson & Heeger (2006)

Now that we have derived two sets of equations for the location function (i.e. with, and without, a constant term in Section 2.1 and 2.3, respectively) let us illustrate the difference with data on the cortical map. The first example are data from Larsson & Heeger (2006, Fig. 5) for area V1. As a reminder, this is about eq. (16) on the one hand – in essence 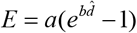, derived from eq. (6) – and the discouraged eq. (18) or (19) on the other hand (in essence 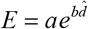, i.e. no constant term outside the exponent – Larsson & Heeger’s constant *‘18.0’ within* the exponent is part of the coefficient *a*).

For the reasons explained above, the retinotopic centre is left undefined by Larsson & Heeger (2006), and a reference eccentricity of *E*_*ref*_ = 3° is used instead. The fitted equation in the original graph in their paper is stated as 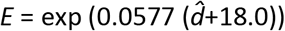, which corresponds to Eq. (18) with constants *a* = 0.0577, and 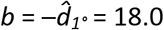. Its graph is shown in Figure 5 as the thick black line copied from the original graph. It is continued to the left as a dotted blue line to show the behaviour toward the retinotopic centre. At the value of 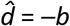, i.e. at a distance of 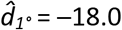 from the 3° representation (as seen from eq. 18 or 19), the line crosses the 1° point. To the left of that point, i.e. towards the retinotopic centre, the curve deviates markedly upward and so the retinotopic centre (*E* = 0°) is never reached.

The pink and the green curve in Figure 5 are two examples for a fit of the equation with a constant term (i.e., for eq. 16). Note that the equations are underconstrained unless either the location of the retinotopic centre or the central CMF *M*_*0*_ are known. The pink curve uses *E*_*2*_ = 0.6° and *d*_*ref*_ = 38 mm, and the green curve *E*_*2*_ = 1.0° and *d*_*ref*_ = 35 mm. Apparently, smaller *E*_*2*_ values go together with larger *d*_*ref*_ values for a similar shape. Within the range of the data set, the two curves fit about equally well; the pink curve is slightly more curved (a smaller *E*_*2*_ is accompanied by more curvature). Below about 1° eccentricity, i.e. around half way between the 3° point and the retinotopic centre, the two curves deviate markedly from the original fit. The new curves fit the data better there than the original and, in particular, reach a retinotopic centre. Of the two, the pink curve (with *E*_*2*_ = 0.6°) reaches the centre at 38 mm from the 3° point, and the green curve at 35 mm.

The centre cortical magnification factor, *M*_*0*_, for the two curves can be derived from eq. (17), giving a value of 35.4 mm/° and 25.3 mm/°, respectively. These two estimates differ substantially from one another – by a factor of 1.4 – even though there is only a 3-mm difference of the assumed location of the retinotopic centre. This illustrates the large effect of the estimate for the centre’s location on the foveal magnification factor, *M*_*0*_. It also illustrates the importance of a good estimate for that location.

There is a graphic interpretation of the foveal magnification factor *M*_*0*_ in these graphs. From eq. (6) one can derive that 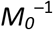 is equal to the function’s slope at the retinotopic centre. Thus, if the function starts more steeply (as does the green curve compared to the pink one), 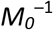 is higher and thus *M*_*0*_ is smaller.

The figure also shows two additional curves (black and brown), depicting data from Duncan & Boynton (2003), which are discussed below. To better display the various curves’ shapes, they are shown again in Figure 6 but now without the data symbols. Figure 6 also includes an additional graph, depicting the exponential function 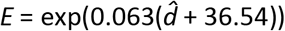 reported by Engel et al. (1994, 1997). In it, 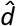 is again the cortical distance in millimetres but this time measured from the 10° representation. *E,* as before, is the visual field eccentricity in degrees. For comparison with the other curves, the curve is shifted (by 19.1 mm cortical distance) on the abscissa, to show the distance from the 3° point. The curve runs closely with that of Larsson & Heeger (2006) and shares its difficulties.

**Figure 6.**
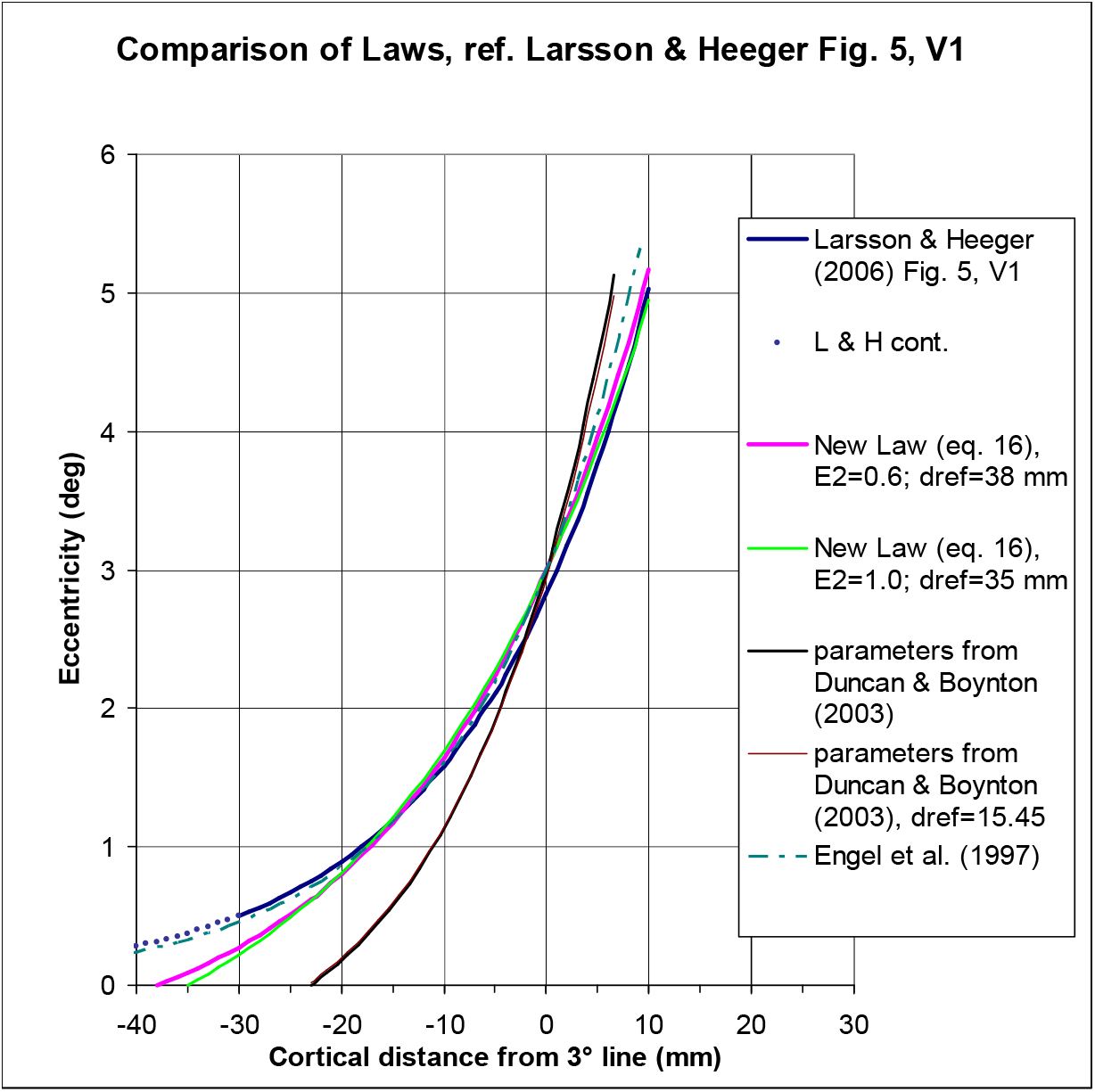
Same as Figure 5 but without the data symbols, for better visibility of the curves. The additional dash-dotted curve next to that of Larsson & Heeger’s depicts the equation by Engel et al. (1997).

#### 2.4.2 The approach of Duncan & Boynton (2003)

In addition to the curves just discussed, Figure 5 and Figure 6 show a further 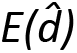 function that is based on the results of Duncan & Boynton (2003). That function obviously differs quite a bit from the others in the figure and it is thus worthwhile studying how Duncan & Boynton (2003) derived these values. The paper takes a somewhat different approach for estimating the retinotopic mapping parameters for V1 than the one discussed before.

As a first step in Duncan & Boynton’s paper, the locations of the lines of equal eccentricity are estimated for five eccentricities (1.5°, 3°, 6°, 9°, 12°) in the central visual field, using the equation *w = k* * log (*z + a*). The function looks similar to the ones discussed above, except that *z* is now a complex variable that mimics the visual field in the complex plane. On the horizontal half-meridian (where *z* is real-valued) that is equivalent to eq. (6) in the present paper, i.e., to an *E(d)* function that includes a constant term (here parameter *a*) in the log’s argument and with the retinotopic centre as the reference. At these locations, the authors then estimate the size of the projection of several 1°-patches of visual space (see their Fig. 3; this is where they differ in their methodology from other approaches). By definition, these sizes are the cortical magnification factors *M*_*i*_ at the corresponding locations. Numerically, these sizes are then plotted vs. eccentricity in the paper’s Fig. 4 (reproduced in Figure 7A). Note that this is not readily apparent from the paper, since both the graph and the accompanying figure caption state something different. In particular the y-axis is labelled incorrectly (as is evident from the accompanying text). For clarity, therefore, Figure 7B here plots these data with a corrected label and on a linear y-axis.

**Figure 7.**
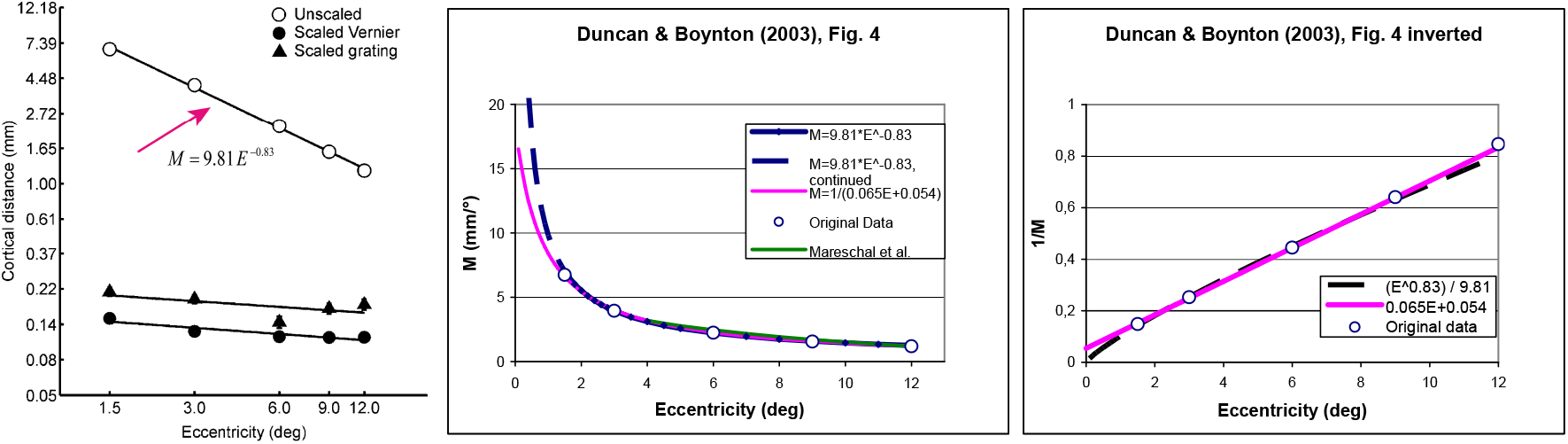
Duncan & Boynton’s (2003) Fig. 4, showing the cortical magnification factor’s variation with eccentricity. (A) Original Fig. 4. The open symbols follow a power function (note the double-linear coordinates). (B) Redrawn on a linear y-axis and with a *corrected* y-axis label (*M* in mm/°). Open circles show the original data. Note that the equation used in (A) and proposed earlier in the paper (p. 662), *M* = 9.81\**E*^−0.83^, predicts an *infinite* foveal magnification factor, shown as the blue curve (with blue diamonds for visibility). In contrast, the inverse-linear fit *M*^−1^ = 0.065 *E* + 0.054 proposed later in the paper (p. 666) fits the data equally well in the measured range of 1.5° to 12° but in contrast predicts a reasonable foveal magnification factor *M*_*0*_ of 18.5 mm/°. The *E*_*2*_ value for the latter equation is *E*_*2*_ = 0.83. The additional green curve shows an equation by Mareschal et al. (2010) (see next section). (C) The inverse of the same functions. Note the slight but important difference at 0° eccentricity, where the original curve is zero and its inverse is thus undefined, whilst the linear function is *non-zero* and its inverse thus well-defined.

The authors next fit a power function to those data, stated as *M* = 9.81\**E* ^−0.83^ for the cortical magnification factor (note the double-logarithmic coordinates in 7A). There is more confusion, however, because it is said that, from such power functions, the foveal value can be derived by extrapolating the fit to the fovea (p. 666). That cannot be the case, however, since, by the definition of a power function (including those used in the paper), there is no constant term. The function therefore goes to infinity towards the fovea centre, as shown in Figure 7B (dashed line). Furthermore, *E*_*2*_, which is said to be derived in this way in the paper, cannot be derived from a nonlinear function (because the *E*_*2*_ concept requires a linear or inverse-linear function). The puzzle is resolved with a reanalysis of Duncan & Boynton’s Fig. 4. It reveals how the foveal value and the connected parameter *E*_*2*_ were, in fact, derived: as an inverse-linear function which fits the data equally well in the measured range of 1.5° – 12° eccentricity (Figure 7B and 7C, continuous line; note the slight but crucial difference in 7C the retinotopic centre). From that function, the foveal value and *E*_*2*_ are readily derived. Indeed, the two values correspond to the values given in the paper.

The distance of the isoeccentricity lines from the retinotopic centre is not specified in Duncan & Boynton (2003). We can derive it from eq. (17), though, because *M*_*0*_ and *E*_*2*_ are fixed:

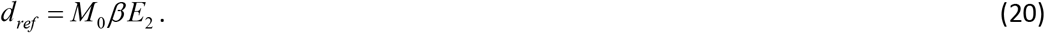

With the authors’ parameters (*M*_*0*_ = 18.5 mm/° and *E*_*2*_ = 0.83), the scaling factor *β* in that equation comes out as *β* = 1.03 (from eq. 16). From that, *d*_*ref*_ = *d*_*1.5°*_ = 15.87 mm. As a further check, we can also derive a direct estimate of *d*_*ref*_ from their Fig. 3. For their subject ROD, for example, the 1.5° line is at a distance of *d*_*1.5°*_ = 15.45 mm on the horizontal meridian. That value is only very slightly smaller than the one derived above. For illustration, Figure 5 and Figure 6 in the previous section also contain a graph for that value (thin black line). Conversely, with *d*_*ref*_ given, *M*_*0*_ can be derived from eq. (17) (or eq. 20), which gives a slightly smaller value of *M*_*0*_ = 18.0 mm/°. The two curves are hardly distinguishable; thus, as previously stated, *d*_*ref*_ and *M*_*0*_ interact, with different value-pairs resulting in similarly good fits.

In summary, the parameters in Duncan & Boynton’s (2003) paper: *M*_*0*_ = 18.5 mm/° and *E*_*2*_ = 0.83, are supported by direct estimates of the size of 1°-projections. They are taken at locations estimated from a set of mapping templates, which themselves are derived from a realistic distance-vs.-eccentricity equation. The paper provides another good example how the linear concept for the magnification function can be brought together with the exponential (or logarithmic) location function. The estimate of *M*_*0*_ comes out considerably lower than in more recent papers (e.g. Schira et al., 2009; see Figure 8 below). Possibly the direct estimation of *M* at small eccentricities is less reliable than the approach taken in those papers.

**Figure 8.**
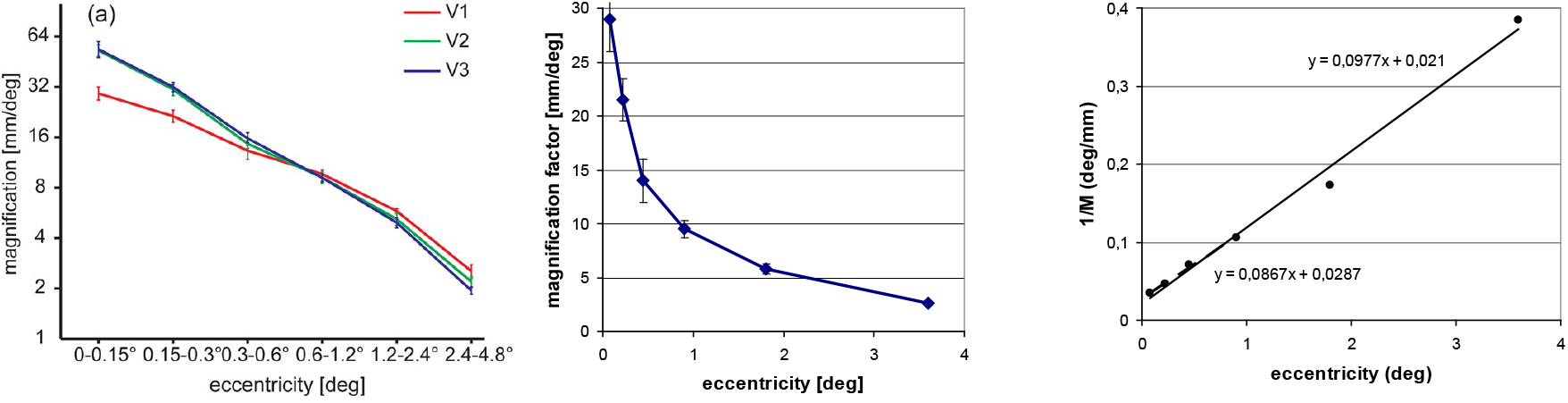
The cortical magnification factor’s dependency on eccentricity from Schira, Tyler, Breakspear & Spehar (2009, Fig. 7A). (A) Original graph. (B) V1 data for *M*, from Schira et al.’s graph but drawn on double-linear coordinates, showing the hyperbola. (C) Resulting inverse factor, again on linear coordinates. The regression line, *M*^−1^ = 0.0977 *E* + 0.021, fits the whole set and predicts *E* = 0.21° and *M*_*0*_ = 47.6 mm. The regression equation *M*^−1^ = 0.0867 *E* + 0.0287 is a fit to only the first four points and might be a better predictor for the retinotopic centre, resulting in the values *E*_*2*_ = 0.33° and *M*_*0*_ = 34.8 mm.

#### 2.4.3 Mareschal, Morgan & Solomon (2010)

Figure 7 shows an additional curve from a paper by Mareschal et al. (2010) on cortical distance, who base their cortical location function partly on the equation of Duncan & Boynton (2003). Mareschal et al. (2010) state their location function as

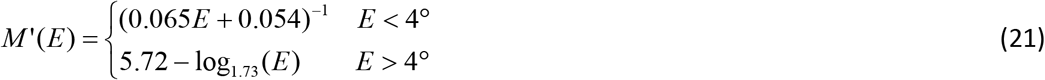

The upper part of the equation is that of Duncan & Boynton (pink curve) and is used below an eccentricity of 4°. The green continuous line shows Mareschal’s log equation above 4°, and the dashed line shows how the log function would continue for values below 4°. As in the previous examples, the latter is not meaningful and is undefined at zero eccentricity, which is why Mareschal et al. switched to the inverse-linear function (i.e. the pink curve) at that point. The problem at low eccentricity is apparent in Fig. 9 of their paper where the x-axis stops at ½ deg, so the anomaly is not fully seen. For their analysis, the switch of functions is not relevant since eccentricities other than 4° and 10° were not tested. However, the example is added here to illustrate that the case distinction in eq. (21) could be avoided with the new equations derived here.

**Figure 9.**
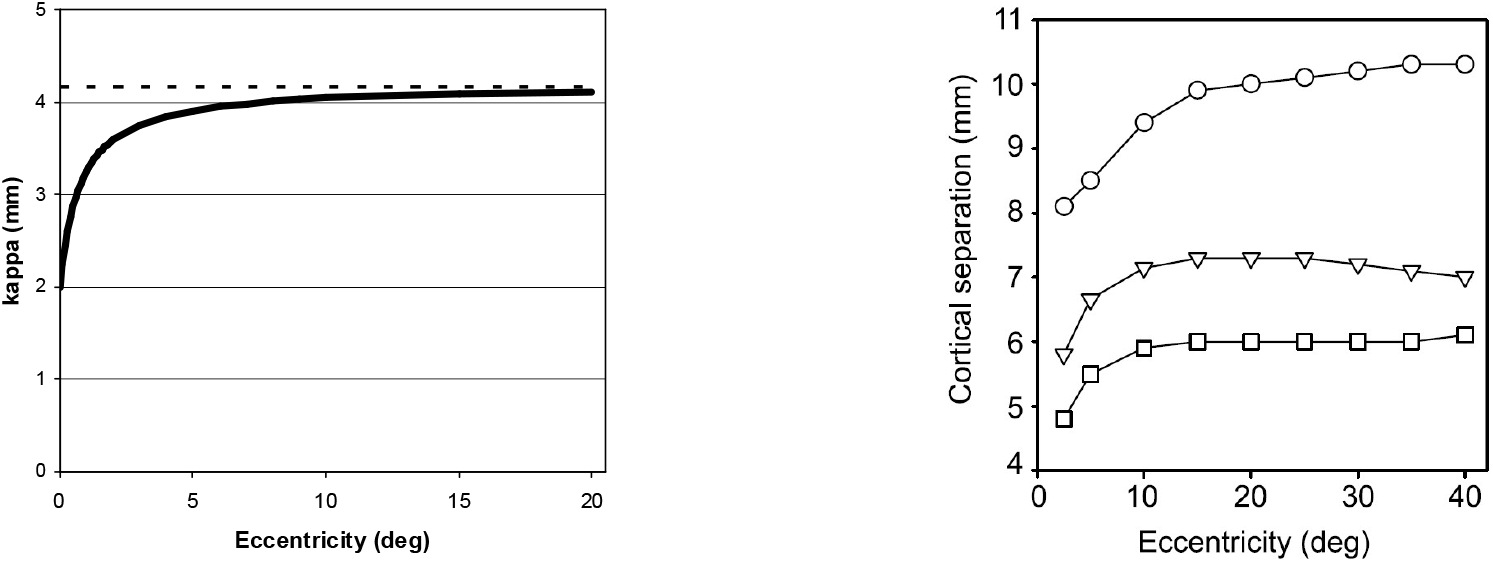
(A) Graph of eq. (31) with realistic values for *M*_*0*_, *E*_*2*_, 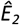, and *δ*_0_. The value of *E*_*2*_ for *M*^*−1*^ was chosen as *E*_*2*_ = 0.8° from Dow, Snyder, Vautin, & Bauer, 1981 (as cited in Levi et al., 1985, or Strasburger et al., 2011, Table 4). *M*_*0*_ = 29.1 mm was chosen to give a good fit with this *E*_*2*_ in Fig. 2. Foveal critical distance was set to *δ*_*0*_ = 0.1° from Siderov, Waugh, & Bedell, 2013, 2014. An 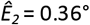 would obtain with this *δ*_*0*_ and the value of *δ*_*4°*_ = 1.2° in Strasburger et al., 1991; it also serves as an example for being a clearly different value than *E*_*2*_ for the cortical magnification factor, to see the influence of the 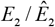 ratio on the graph. Cortical critical distance *κ* starts from the value given in eq. (32) for the fovea centre (around 2 mm) and converges to the value in eq. (33). Cortical critical distance for crowding from Motter & Simoni (2007, Fig. 7), showing the qualitative similarity for the dependency. The curves are effectively based on Duncan & Boynton’s (2003) inverse-linear equation (see Figure 7B above, pink curve), which implies *M*_*0*_ = 18.5 mm/° und *E*_*2*_ = 0.83°. The middle curve (triangles) is comparable to the curve in (A). The different asymptote in (B) stems from a different *M*_*0*_.

#### 2.4.4 An added exponent: Sereno et al. (1995)

To accommodate for a slight curvature in the inverse CMF function (Figure 2A), several authors have suggested using a modestly nonlinear function for its modelling (Rovamo & Virsu, 1979; Van Essen et al., 1984; cf. Table 1). One way to achieve this is using a power function, i.e., adding an exponent to the linear function with a value slightly above 1:

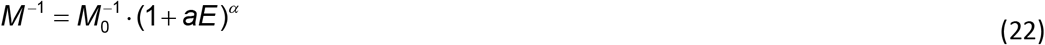

Van Essen et al. (1984), e.g., use an exponent of 1.1. Following their lead, Sereno et al. ((1995)) posit

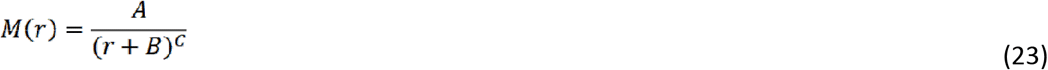

for the CMF, where *A, B,* and exponent *C* are free parameters, and *r* denotes eccentricity along a radius (the equations are found in the paper’s footnotes 24, 25, and 26). For the case *C* = 0, the equation is reduced to the standard inverse-linear function (eq. 1). By integration, they derive from that the cortical location function, called *mapping function D(r)* = ∫ *M(r)dr* in their paper:

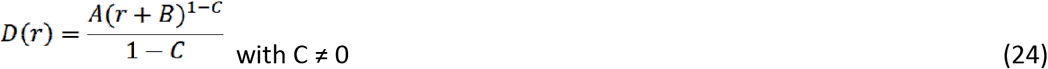

In their fits to the anatomical data, *C* comes out with values close to 1.

Note that both eq. (22) and (23) are well-defined and meaningful in the retinotopic centre (*r*=0). Note also, however, that the exponent *(C)* must not be zero for the location function (eq. 24). I.e., the location function is undefined for the inverse-linear CMF function. That latter case is discussed in Sereno et al.’s Footnote 26, where *C* = 1; the cortical location function is then said to converge to

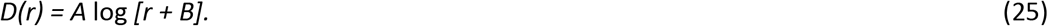

(i.e., similar to eq. 5).

In that equation, however, lies the fatal error that led to the avoidance of the (much simpler) logarithmic location function. On closer inspection and comparison to eq. (5), one can see that, even though there is a constant term (namely *B*), the scaling factor for the independent variable *r* is missing. The equation should be something like *D(r)* = *A* log [***C****r + B]*. Therefore, *B* is effectively constrained to 1 because only then is *D(r*=0) = 0. In other words, the constant term *B* is not actually a free parameter.

Interestingly, Sereno et al. (1995) are aware of the shortcomings of the latter equation. They write, “Our data could also be fit with this equation, but only if we allowed B to be negative, which results in a singularity (infinite magnification factor) before the center-of-gaze is reached. A good fit without a singularity could only be achieved with *C* above 1”. They continue saying, “The combinations of parameters given here fit the cortical distance data [i.e., referring to the location function] very closely but still give unrealistically large estimates of cortical magnification at the exact center of the fovea […], indicating that the standard equation for M is inadequate to accurately describe cortical magnification in the very center of the fovea in humans even with C > 1.” What went unnoticed is that a simple remedy would have been using the correct additive constant term and a scaling factor for *r*.

#### 2.4.5 Toward the retinotopic centre: Schira et al.

As discussed above, predictions on the properties at the retinotopic centre depend critically on determining its precise location and thus require data at small eccentricities. Schira, Tyler and coworkers have addressed that problem in a series of papers (Schira et al., 2007; Schira, Tyler, Breakspear, & Spehar, 2009; Schira et al., 2010) and provide detailed maps of the centres of the early visual areas, down to 0.075° eccentricity. They also develop parametric, closed analytical equations for the 2D maps. When considered for the horizontal direction only, these equations correspond to those discussed above (eq. 1 and eq. 16/17) (the equations differ on, and close to, the vertical meridian – Schira et al., 2007; Schira et al., 2010 – but this is not relevant here).

Figure 8 shows magnification factors from Schira et al., 2009, Fig. 7A, with figure part B showing their V1 data (red curve), but redrawn on double-linear coordinates. As can be seen, the curve runs close to a hyperbola. Its inverse is shown in Figure 8C, which displays the familiar, close-to-linear behaviour over a wide range, with a positive y-axis intercept that corresponds to the value at the retinotopic centre, 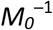. From the regression line, *M*_*0*_ and *E*_*2*_ are readily obtained and are *E*_*2*_ = 0.21° and *M*_*0*_ = 47.6 mm, respectively. Interestingly, a rather large value of *M*_*0*_ is obtained compared to previous reports. Partly (as can also be seen from the graph) that can be caused by a single, most peripheral point; the centrally located values predict a somewhat shallower slope of the linear function. If one disregards that point in the regression, one arrives at a slightly larger *E*_*2*_ and smaller *M*_*0*_ value: *E*_*2*_ = 0.33° and *M*_*0*_ = 34.8 mm. The latter values might be the more accurate predictors for V1’s central point.

In summary, the derived equations provide a direct link between the nomenclature more well-known in psychophysics and that in the neurophysiological literature on retinotopy. They were applied to data for V1 (Fig. 2) but will work equally well for higher early visual areas, including V2, V3, and V4 (cf. Larsson & Heeger, 2006, Fig. 5; Schira et al., 2009, Fig. 7). *M*_*0*_ is expected to be slightly different for the other areas (Schira et al., 2009, Fig. 7) and so will likely be the other parameters.

#### 2.4.6 *d*_*2*_ – a structural parameter to describe the cortical map

As shown in Section 2.1, a newly defined structural parameter *d*_*2*_ can be used to describe the cortical location function very concisely (eq. 9 or 10). Parameter *d*_*2*_ is the cortical representation of Levi and Klein’s *E*_*2*_. That is, *d*_*2*_ is the distance from the retinotopic centre, measured in mm, corresponding to eccentricity *E*_*2*_, which is where the foveal value doubles. Eq. (8) can serve as a means to obtain an estimate for *d*_*2*_. Essentially, *d*_*2*_ is the product of *M*_*0*_ and *E*_*2*_ with a scaling factor. Table 2 gives a summary of *d*_*2*_ estimates thus derived. The value of *d*_*2*_ ≈ 8 mm with *E*_*2*_ = 0.33°, based on Schira et al.’s (2009) data which go down to very low eccentricities, might be the most accurate estimate currently given their sophisticated methodology for assessing the map closely around the retinotopic centre.

Similar to what *E*_*2*_ does for the linear or inverse-linear function – be it the anatomical CMF or thresholds in a psychophysical task – *d*_*2*_ concisely captures the properties of the map in a single number. It is given in physical units (mm) and can thus be drawn directly into a retinotopic map. *E*_*2*_ can be (and has been) used as a summary measure for the CMF but is not as well-suited because its units are in deg visual angle on the retina (or in the visual field), i.e. needs to be translated to spatial, cortical units. Currently, typical characterisations of the cortical map are done by drawing iso-eccentricity lines at several eccentricities (10°, 20°, 30°, etc.). In a similar way, a single *d*_*2*_ line could be drawn on the cortical map, or *d*_*2*_ could marked as a point on a radius. As a characteristic measure, *d*_*2*_ could be used in many ways, for comparison of the anisotropy in the cortical maps, between species, individuals, gender, etc. Or indeed it could describe any other retinotopic map like those for V2 – V4 or that for the LGN, the pulvinar, the reticular nucleus of the thalamus, once data are available. Differences between *d*_*2*_ show a difference in the architecture.

That said, *d*_*2*_ shares certain limitations of *E*_*2*_ (Strasburger et al., 2011, Fig. 11 and Table 3). Like the latter, it relies on data in, and near, the retinotopic centre and can thus be expected to be most meaningful at small to medium eccentricities. Its validity for describing the curve at larger eccentricities further depends on the premise that location in the map results from integrating the local magnification function, i.e., that local magnification factors “add up”. For the CMF, that appears to be the case, as evidenced by the good fit of location data shown in Figure 5 and other log location functions in the literature. Yet for local properties that are likely based on differences in neural wiring, like the colour channels studies in D’Souza et al. (2016), that might not be the case. *d*_*2*_, in those cases, will characterise the function, but not its map.

Note in Table 2 that both the estimates for *M*_*0*_ (the central CMF for V1) and *E*_*2*_ vary quite a bit between neuroanatomical studies. Except for Dougherty et al.’s (2003, Fig. 5) estimate, current *M*_*0*_ estimates are much larger than the old estimates of *M*_*0*_ = 8.55 mm/° from Cowey & Rolls (1974) or *M*_*0*_ = 7.99 mm/° from Rovamo & Virsu (1979) (for more estimates of *M*_*0*_, see Strasburger et al., 2011, Table 5). At the same time, again except Dougherty et al. (2003, Fig. 5), modern *E*_*2*_ values for the CMF on the whole appear smaller than the old values (Cowey & Rolls: 1.75°, Rovamo & Virsu: 3.0°). Since, in essence *d*_*2*_ is the product of the two, these variations in opposite directions are evened by *d*_*2*_ which indeed varies less between studies. This might be another reason why *d*_*2*_ could be a more suitable structural parameter for a retinotopic map than either *M*_*0*_ or *E*_*2*_ on its own.

**Table 2.**
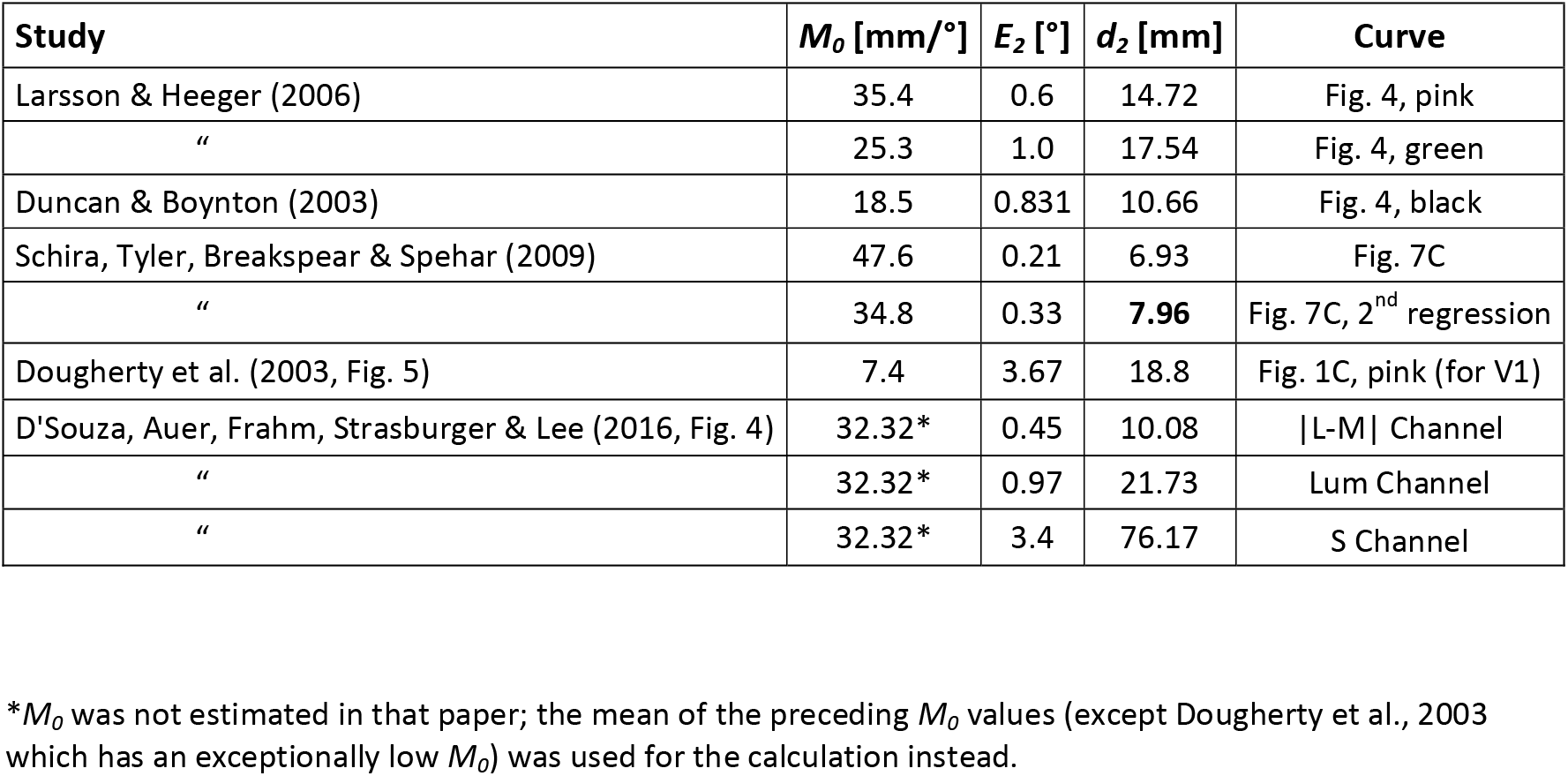
Values of the parameter *d*_*2*_ from an analysis of data in several studies, by eq. (8): *d*_*2*_ = *M*_*0*_ *E*_*2*_ ln(2). *d*_*2*_ is the cortical representation of *E*_*2*_ and characterizes the cortical location function in a single value.

## 3. Crowding and Bouma’s Law in the cortex

The preceding sections were about the cortical location function; in the final section that function will be applied to an important property of cortical organization: visual crowding. Whereas, in the preceding, cortical *location* was the target of interest, in this section we are concerned with cortical *distances*.

As reviewed in the introduction, MAR-like functions like acuity generally change in peripheral vision in that *critical size* scales with eccentricity, so deficits can (mostly) be compensated for by *M*-scaling (as, e.g. in Rovamo & Virsu, 1979). For crowding, in contrast, target size plays little role (Strasburger et al., 1991; Pelli et al., 2004; Whitney & Levi, 2011). Instead, the critical *distance* between target and flankers scales with eccentricity, though at a different rate than MAR (Rosenholtz, 2016; Strasburger, 2020). This scaling characteristic of crowding is known as Bouma’s rule or Bouma’s law (Bouma, 1970; Strasburger et al., 1991; Pelli et al., 2004; Pelli & Tillman, 2008; Strasburger, 2020). The corresponding distances in the primary cortical map are thus governed by *differences* of the cortical location function as derived here in Section 2. Crowding’s critical distance (or indeed any distance, including acuity gap size) is thus, in a sense, a spatial derivative of location. Pattern recognition, at even slight eccentricities, is governed by the crowding phenomenon and is largely unrelated to visual acuity (or thus to cortical magnification) (Strasburger et al., 1991; Pelli et al., 2004; Pelli et al., 2007; Pelli & Tillman, 2008; Strasburger & Wade, 2015). For understanding crowding it is paramount to look at its cortical basis, since we know since Flom, Weymouth, & Kahnemann (1963) that crowding is of cortical origin (as also emphasized by Pelli, 2008).

A question that arises naturally in that context then is how the cortical equivalent of critical crowding distance varies across the visual field. Klein & Levi (1987) were the first to consider a related question, namely how the cortical distance for distance threshold in a vernier task varies with eccentricity. They conclude that it is approximately constant. That conclusion was based on the observation that taking the first derivative of Schwartz’s (1980) log mapping using the constancy assumption will result in the well-known inverse-linear cortical magnification function. Conversely, their empirically determined position thresholds, when mapped by an inverse-linear cortical magnification function (with an *E*_*2*_ of 0.6), turned out mostly constant across a wide range of eccentricities (cf. Klein & Levi’s Fig. 5). Later, Duncan and Boynton (2003), after estimating *M* based on Schwartz’s (1980) log mapping and applying that to obtain cortical distances (see Section 2.4.2), show that, for scaled vernier tasks and scaled gratings, the cortical equivalents are again mostly constant (above 1.5° eccentricity; 2003, Fig. 4). Similarly, with respect to the cortical distance for crowding’s critical distance, it has been proposed that it is likely a constant, with the same reasoning Motter & Simoni, 2007; Pelli, 2008; Mareschal, Morgan, & Solomon, 2010; oddly, the original source for the log mapping, Fischer, 1973, is not cited in the above papers).

Elegant as it seems as a take-home message, however, the constancy assumption is most likely incorrect as a general rule and is only true at sufficiently large eccentricities. If stated as a general rule, it rests on the same shortcut of equating linearity and proportionality, i. e. the omission of the constant term that gave rise to those cortical location functions that miss the retinotopic centre (Section 2.3). Based on the properties of the cortical location function derived in Section 2, it will turn out that the critical cortical crowding distance increases steeply within the fovea (where, e.g., reading mostly takes place) and reaches an asymptote beyond perhaps 5° eccentricity, consistent with a constancy at sufficient eccentricity. Accordingly, Pelli (2008) warns against extrapolating the constancy toward the retinotopic centre. Remarkably (and to my pleasant surprise), after I had completed the derivations it turned out that the analytic equation exposed below nicely agrees with those presented by Motter & Simoni (2007, Fig. 7). In that figure, reproduced here in Figure 9B, only the more peripheral data above about 10° show the presumed constancy.

Let us turn to the equations. Bouma (1970) stated what is now known as Bouma’s law for crowding (Strasburger, 2020):

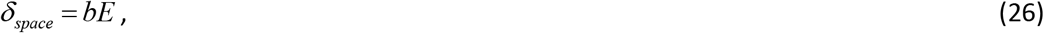

where *δ*_*space*_ is the free space between the patterns at the critical distance and *b* is a proportionality factor. Bouma (1970) proposed an approximate value of *b* = 0.5 = 50%, which is now widely cited, but he also mentioned that other proportionality factors might work equally well; indeed, Pelli et al. (2004) have shown that *b* can take quite different values, depending on the exact visual task. Yet even though the factor may be different between tasks, the implied linearity of eq. (26) almost always holds up. The law could thus be restated as saying that free space for critical spacing is proportional to eccentricity, with the proportionality factor taking some value around 50% or 40%, depending on the task.

In today’ literature it has become customary to state flanker distance not as free space but as measured from the respective centres of target and flankers. The critical spacing then remains largely constant across sizes as Tripathy & Cavanagh, 2002 and others have shown. To restate Bouma’s law for that centre-to-centre distance *δ*, let the target pattern have a size *S* in the radial direction (e.g., *width* in the horizontal), so that *δ* = *S* + *δ*_*space*_. Then eq. (26) becomes

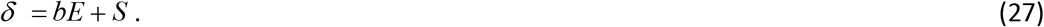

This equation no longer represents proportionality yet is still linear in *E*. Importantly, however, going from Bouma’s equation (eq. 26) to that in eq. (27) reflects adding the constant term that we talked about in the preceding sections. And formally, that equation (27) is analogous to size scaling as in. (2). Analogously to Levi and Klein’s *E*_*2*_ we therefore introduce a parameter 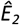 for crowding, as *the* eccentricity where the *foveal value of critical distance doubles*. Denoting the foveal value of critical distance by *δ*_0_, we get, from eq. (27):

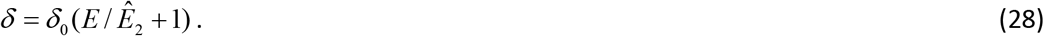

Obviously, that equation is analogous to eq. (1) and (2) that we started out with; it describes how critical distance in crowding is linearly dependent on (but is not proportional to) eccentricity in the visual field. In this respect, it thus behaves like acuity and many other spatial visual performance measures, just with a different slope and axis intercept.

With the equations derived in the preceding sections, we can now derive the critical crowding distance in the cortical map, i.e. the cortical representation of critical distance in the visual field. Let us denote that distance by κ (kappa). By definition, it is the difference between the map locations for the target and a flanker at the critical distance in the crowding task: *κ* = *d*_*f*_ − *d*_*t*_. The two cortical locations *d*_*f*_ and *d*_*t*_ are, in turn, obtained from the mapping function, which is given by inverting eq. (6) above:

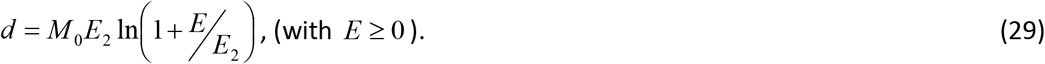

As before, *d* is the distance of the location in the cortical map from the retinotopic centre. So, critical distance κ for crowding in the retinotopic map is the difference of the respective *d* values for target and flanker, *κ* = *d*_*f*_ − *d*_*t*_ :

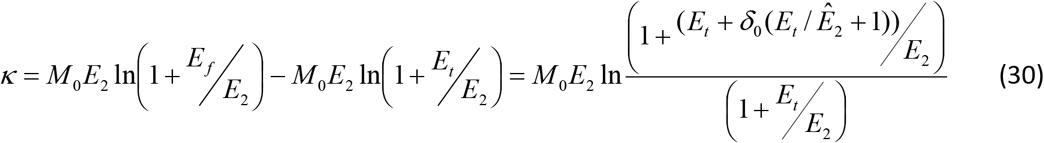

(by eq. 29 and 28), where *E*_*t*_ and *E*_*f*_ are the respective eccentricities at which target and flanker are located.

After simplifying and setting target eccentricity *E*_*t*_ = *E* for generality, this becomes

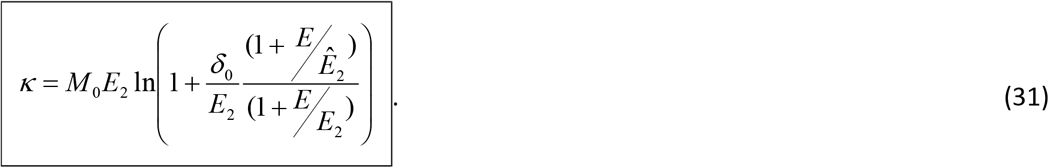

Note that we stated that equation previously (Strasburger & Malania, 2013, eq. 13, and Strasburger et al., 2011, eq. 28), but, alas, incorrectly: a factor was missing.

To explore this function, its graph is shown in Figure 9A and we look at two special cases. In the retinotopic centre, equation (31) predicts a critical distance *κ*_0_ in the cortical map of

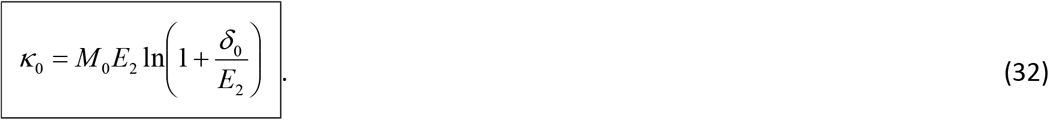

With increasing eccentricity, κ departs from that foveal value and increases, depending on the ratio 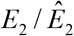 (provided 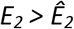 which can be reasonably assumed; Latham & Whittaker (1996; Strasburger, 2020). Numerator and denominator are the *E*_*2*_ values for the location function and the crowding function, respectively (eq. 1 vs. eq. 28). They are generally different, so their ratio is not unity.

With sufficiently large eccentricity, the equation converges to

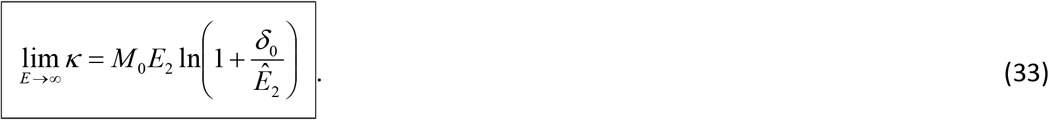

The expression is shown as dashed line in Figure 9A. It is identical to that for the foveal value in eq. (33) except that *E*_*2*_ is now replaced by the corresponding value 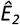 for crowding.

Importantly, note that *kappa* varies substantially around the centre, by around two-fold between the centre and 5° eccentricity with realistic values of *E*_*2*_ and 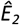. This, as said above, is at odds with the conjecture that the cortical critical crowding distance is basically a constant (Motter & Simoni, 2007; Pelli, 2008; Mareschal et al., 2010). Pelli (2008) presented a mathematical derivation for the constancy, very similar to the one presented above – based on Bouma’s law and Schwartz’ (1980) logarithmic mapping function. The discrepancy arises from the underlying assumptions: Pelli used Bouma’s law as proportionality, i.e., in its simplified form stated in eq. (26) (its graph passing through the origin). The simplification was done on the grounds that the error is small outside the retinotopic centre and plays little role; the paper appropriately warns that additional provisions must be made at small eccentricities. Schwartz’s (1980) (simplified) mapping function was consequently also used in its simplified form (without the constant term), for the same reason. With these simplifications the critical distance in the cortex indeed turns out as simply a constant.

As should be expected, at sufficiently high eccentricities κ is close to constant in the derivations given above (Figure 9). These equations (eq. 31–33) can thus be seen as a generalization of Pelli’s result that now also covers the (obviously important) case of central vision.

For comparison, Figure 9B shows critical crowding spacing on the cortical map from a paper on visual search by Motter & Simoni (2007). Note that the shown curves, though inspired by their experimental search data, are not based on these but are based on a cortical-surface model (shown in their Fig. 1), obtained by *M*-scaling visual distances. Critical distances are assumed to follow Bouma’s law, with a Bouma factor of ½. *M*-scaling is by Duncan & Boynton’s (2003) inverse-linear equation (1/*M(w)* = 0.065*w* + 0.054; shown here in Figure 7B above, pink curve). The figure’s basis is thus the same as in the present paper and effectively shows Bouma’s law mapped onto the cortex. The three curves refer to different flanker location and reflect crowding asymmetry (see Strasburger, 2020, for review) (upper curve: peripheral flanker, lower curve: central flanker); the middle curve is is for equal-eccentricity flanker distances and is the one comparable to the curve in (A). Duncan & Boynton’s equation implies *M*_*0*_ = 18.5 mm/° and *E*_*2*_ = 0.83° (cf. Table 2 above). That *E*_*2*_ is similar to that assumed in Figure 9A; *M*_*0*_ is different. As we have seen in eq. (33), the asymptote depends on these two values. The different asymptote in 9B thus stems from the different *M*_*0*_.

Pelli & Tillman (2008, Online Supplement) derive a value of 6 mm for the asymptote. It is based on eq. (18) above, as reported by Larsson & Heeger (2006), and a Bouma factor of 0.4.

An interesting (though unrealistic) special case of eq. (31) is the one in which *E*_*2*_ and 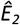 are equal. *κ* is then a constant, as Pelli (2008) predicted. Its value in that case would be simply given by

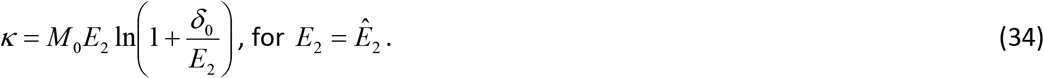

On a different note, equations (31)–(34) have *M*_*0*_ as a scaling factor and, as said before, *M*_*0*_ appears to be more difficult to determine empirically. However, *M*_*0*_ can be replaced, as shown above. From eq. (17) we know that

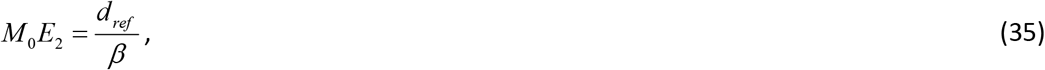

which, by the definition of *β*, takes a particularly simple form when we choose *d*_*2*_ (the cortical equivalent of *E*_*2*_) as the reference:

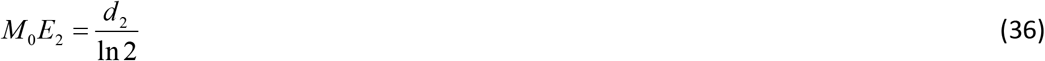

(this is the same as eq. 8a). We can then rewrite the equation for the critical cortical crowding distance (eq. 31) as

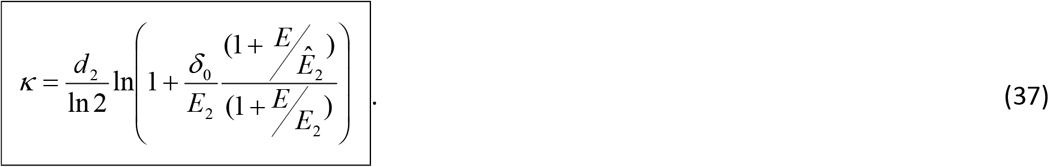

Similarly, the two special cases given in eq. (32) and (33) become

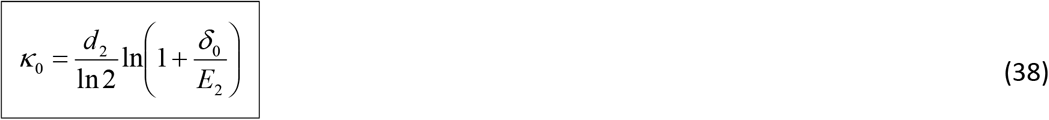

and

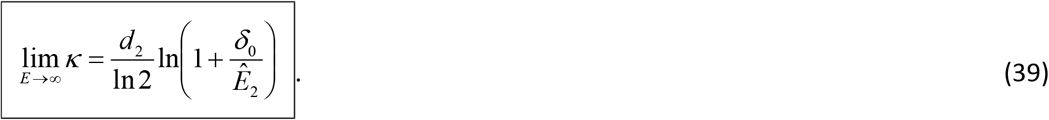

Values for *d*_*2*_ derived from the literature by eq. (36) that could be plugged into eq. (38) and (39) *were provided in Table 2 above*. These two equations ((38) and (39)), for the retinotopic centre and eccentricities above around 5°, respectively, could lend themselves for determining critical crowding distance in the cortex.

In summary for the cortical crowding distance, the two well-established linear eccentricity laws –for cortical magnification in neuroscience and critical crowding distance in psychophysics –together with Fischer’s (1973) or Schwartz’s (1977; 1980) equally well-established logarithmic mapping rule, predict a highly systematic behaviour of crowding’s critical distance in the cortical map. Given the very similar mappings in areas V2, V3, V4 (Larsson & Heeger, 2006; Schira et al., 2009), that relationship can be expected to be similar in those areas as well (see Figure 9A for a graph). Since the equations for crowding follow mathematically, they should work well there with suitable *E*_*2*_ values inserted. Thus, direct confirmations of their behaviour can cross-validate mapping models and might shed light on the cortical mechanisms underlying crowding.

## 4. Outlook

Where does this leave us? The early cortical visual areas are very regularly organized and their spatial maps appear to be pretty similar. Yet variation of perceptual performance across the visual field differs widely between visual tasks, as highlighted by their respective, widely differing *E*_*2*_ values. For cortical magnification, in contrast, *E*_*2*_ estimates appear quite similar to each other. It is not yet clear how different spatial scalings in psychophysics can emerge from a largely uniform cortical architecture when there can be only one valid location function on any radius. The equivalence between psychophysical *E*_*2*_ and the cortical location function in the preceding equations thus likely only hold for a single *E*_*2*_, presumably the one pertaining to low-level tasks that are somehow connected to stimulus size. 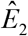 for critical crowding distance would be an example for a psychophysical descriptor that is decidedly not related to stimulus size (Tripathy & Cavanagh, 2002; Pelli et al., 2004); it rather reflects location differences. The underlying cortical architecture that brings about psychophysical *E*_*2*_ values different from that of the CMF (like 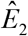) could be neural wiring differences, within or between early visual areas, underneath a similar topography.

The link between the (local) CMF function and the (global) cortical location function derived here rests on the assumption of spatial additivity – that local distances add-up to global distances and the location function is thus the integral of the CMF function. *E*_*2*_ values different from that of the CMF thus do not translate to a location function. When two different *E*_*2*_ values act together, as in crowding, nonlinear functions as those in Figure 9 arise.

To go further, one of the basic messages of the cortical-magnification literature is the realization that by *M*-scaling stimulus size some, but not all, performance variations are equalised across the visual field. In parameter space, these other variables can be said to be *orthogonal* to target size. Pattern contrast is such a variable (Strasburger, Rentschler, & Harvey, 1994) which needs to be scaled independently from size to equalize performance in pattern recognition. Temporal resolution is another example (Poggel, Calmanti, Treutwein, & Strasburger, 2012). Again, differing patterns of connectivity between retinal cell types, visual areas, and along different processing streams likely underlie these performance differences. The aim of the present paper is just to point out that a *common spatial location function* underlies the early cortical architecture that can be described by a unified equation. This equation includes the fovea including the retinotopic centre, and has parameters that are common in psychophysics and physiology.

## Acknowledgements

I thank Dany d’Souza for the original question, Barry Lee for critical comments on the manuscript and meticulous language corrections, Zhaoping Li and Josh Solomon for critical reading, and Zhaoping Li for a thorough check of the mathematical derivations. Thanks also to an anonymous reviewer for pointing out the paper by Klein & Levi (1987) and a reviewer who pointed out misleading phrasings.

## Footnotes

1 Note that the elegance of the complex-log representation is deceiving in that not all properties of the complex plane have a counterpart in the 2D real plane (which is undesirable for a mathematical representation). For example, the square of a value on the upper vertical meridian does not correspond to a value on the left horizontal meridian, as would be implied by i^2^ = −1.

## References

Aubert, H. R., & Foerster, C. F. R. (1857). Beiträge zur Kenntniss des indirecten Sehens. (I). Untersuchungen über den Raumsinn der Retina. Archiv für Ophthalmologie, 3, 1–37.

Bouma, H. (1970). Interaction effects in parafoveal letter recognition. Nature, 226, 177–178.

Cowey, A., & Rolls, E. T. (1974). Human cortical magnification factor and its relation to visual acuity. Experimental Brain Research, 21, 447–454.

D’Souza, D. V., Auer, T., Strasburger, H., Frahm, J., & Lee, B. B. (2016). Dependence of chromatic response in V1 on visual field eccentricity and spatial frequency: an fMRI study. Journal of the Optical Society of America A, 33(3), A53–A64.

Daniel, P. M., & Whitteridge, D. (1961). The representation of the visual field on the cerebral cortex in monkeys. Journal of Physiology, 159, 203–221.

Dougherty, R. F., Koch, V. M., Brewer, A. A., Fischer, B., Modersitzki, J., & Wandell, B. A. (2003). Visual field representations and locations of visual areas V1/2/3 in human visual cortex. Journal of Vision, 3(10), 586–598.

Dow, B. M., Snyder, R. G., Vautin, R. G., & Bauer, R. (1981). Magnification factor and receptive field size in foveal striate cortex of the monkey. Experimental Brain Research, 44, 213–228.

Duncan, R. O., & Boynton, G. M. (2003). Cortical magnification within human primary visual cortex. Correlates with acuity thresholds. Neuron, 38, 659–671.

Engel, S. A., Glover, G. H., & Wandell, B. A. (1997). Retinotopic organization in human visual cortex and the spatial precision of functional MRI. Cerebral Cortex, 7, 181–192.

Engel, S. A., Rumelhart, D. E., Wandell, B. A., Lee, A. T. L., Glover, G. H., Chichilnisky, E.-J., et al. (1994). fMRI of human visual cortex. Nature, 369(6481), 525.

Fischer, B. (1973). Overlap of receptive field centers and representation of the visual field in the cat’s optic tract. Vision Research, 13, 2113–2120.

Flom, M. C., Weymouth, F. W., & Kahnemann, D. (1963). Visual resolution and contour interaction. J. Opt. Soc. Am. 53, 1026–1032.

Foster, D. H., Thorson, J., McIlwain, J. T., & Biederman-Thorson, M. (1981). The fine-grain movement illusion: A perceptual probe of neural connectivity in the human visual system. Vision Research, 21, 1123–1128.

Harvey, B. M., & Dumoulin, S. O. (2011). The relationship between cortical magnification factor and population receptive field size in human visual cortex: Constancies in cortical architecture. The Journal of Neuroscience, 31(38), 13604–13612.

Harvey, L. O.Jr.,, & Pöppel, E. (1972). Contrast sensitivity of the human retina. American Journal of Optometry and Archives of the American Academy of Optometry, 49, 748–753.

Horton, J. C., & Hoyt, W. F. (1991). The representation of the visual field in human striate cortex. A revision of the classic Holmes map. Archives of Ophthalmology, 109(6), 816–824.

Klein, S. A., & Levi, D. M. (1987). Position sense of the peripheral retina. Journal of the Optical Society of America A, 4(8), 1543–1553.

Larsson, J., & Heeger, D. J. (2006). Two retinotopic visual areas in human lateral occipital cortex. The Journal of Neuroscience, 26(51), 13128–13142.

Latham, K., & Whitaker, D. (1996). Relative roles of resolution and spatial interference in foveal and peripheral vision. Ophthalmic and Physiological Optics, 16, 49–57.

Levi, D. M., Klein, S. A., & Aitsebaomo, A. P. (1984). Detection and discrimination of the direction of motion in central and peripheral vision of normal and amblyopic observers. Vision Research, 24, 789–800.

Levi, D. M., Klein, S. A., & Aitsebaomo, A. P. (1985). Vernier acuity, crowding and cortical magnification. Vision Research, 25, 963–977.

Mäkelä, P., Näsänen, R., Rovamo, J., & Melmoth, D. (2001). Identification of facial images in peripheral vision. Vision Research, 41, 599–610.

Mareschal, I., Morgan, M. J., & Solomon, J. A. (2010). Cortical distance determines whether flankers cause crowding or the tilt illusion. Journal of Vision, 10(8), 13:11–14.

Motter, B. C., & Simoni, D. A. (2007). The roles of cortical image separation and size in active visual search performance. Journal of Vision, 7(2):6, 1–15.

Nandy, A. S., & Tjan, B. S. (2012). Saccade-confounded image statistics explain visual crowding. Nature Neuroscience, 15(3), 463–471.

Oehler, R. (1985). Spatial interactions in the rhesus monkey retina: a behavioural study using the Westheimer paradigm. Experimental Brain Research, 59, 217–225.

Osterberg, G. (1935). Topography of the layer of rods and cones in the human retina. Acta Ophthalmologica. Supplement, 6-10, 11–96.

Pelli, D. G. (2008). Crowding: a cortical constraint on object recognition. Current Opinion in Neurobiology, 18, 445–451.

Pelli, D. G., Palomares, M., & Majaj, N. J. (2004). Crowding is unlike ordinary masking: Distinguishing feature integration from detection. Journal of Vision, 4(12), 1136–1169.

Pelli, D. G., & Tillman, K. A. (2008). The uncrowded window of object recognition. Nature Neuroscience, 11(10), 1129–1135 (plus online supplement).

Pelli, D. G., Tillman, K. A., Freeman, J., Su, M., Berger, T. D., & Majaj, N. J. (2007). Crowding and eccentricity determine reading rate. Journal of Vision, 7(2).

Poggel, D. A., Calmanti, C., Treutwein, B., & Strasburger, H. (2012). The Tölz Temporal Topography Study: Mapping the visual field across the life span. Part II: Cognitive factors shaping visual field maps. Attention, Perception & Psychophysics, 74, 1133–1144.

Pöppel, E., & Harvey, L. O.Jr., (1973). Light-difference threshold and subjective brightness in the periphery of the visual field. Psychologische Forschung, 36, 145–161.

Rovamo, J., & Virsu, V. (1979). An estimation and application of the human cortical magnification factor. Experimental Brain Research, 37, 495–510.

Rovamo, J., Virsu, V., & Näsänen, R. (1978). Cortical magnification factor predicts the photopic contrast sensitivity of peripheral vision. Nature, 271, 54–56.

Schira, M. M., Tyler, C. W., Breakspear, M., & Spehar, B. (2009). The foveal confluence in human visual cortex. The Journal of Neuroscience, 29 (July 15), 9050–9058.

Schira, M. M., Tyler, C. W., Spehar, B., & Breakspear, M. (2010). Modeling magnification and anisotropy in the primate foveal confluence. PLoS Computational Biology, 6(1), e1000651.

Schira, M. M., Wade, A. R., & Tyler, C. W. (2007). Two-dimensional mapping of the central and parafoveal visual field to human visual cortex. Journal of Neurophysiology [Epub ahead of print], 97(6), 4284–4295.

Schwartz, E. L. (1977). Spatial mapping in the primate sensory projection: Analytic structure and relevance to perception. Biological Cybernetics, 25, 181–194.

Schwartz, E. L. (1980). Computational anatomy and functional architecture of striate cortex: A spatial mapping approach to perceptual coding. Vision Research, 20, 645–669.

Sereno, M. I., Dale, A. M., Reppas, J. B., Kwong, K. K., Belliveau, J. W., Brady, T. J., et al. (1995). Borders of multiple visual areas in humans revealed by functional magnetic resonance imaging. Science 268, 889–893.

Siderov, J., Waugh, S. J., & Bedell, H. E. (2013). Foveal contour interaction for low contrast acuity targets. Vision Research, 77, 10–13.

Siderov, J., Waugh, S. J., & Bedell, H. E. (2014). Foveal contour interaction on the edge: response to ‘letter-to-the-editor’ by Drs. Coates and Levi. Vision Research, 96, 145–148.

Slotnick, S. D., Klein, S. A., Carney, T., & Sutter, E. E. (2001). Electrophysiological estimate of human cortical magnification. Clinical Neurophysiology, 112(7), 1349–1356.

Strasburger, H. (2020). Seven myths on crowding and peripheral vision. i-Perception, 11(3), 1–46.

Strasburger, H., Harvey, L. O. J., & Rentschler, I. (1991). Contrast thresholds for identification of numeric characters in direct and eccentric view. Perception & Psychophysics, 49, 495–508.

Strasburger, H., & Malania, M. (2013). Source confusion is a major cause of crowding. Journal of Vision, 13(1), 1–20.

Strasburger, H., Rentschler, I., & Harvey, L. O.Jr., (1994). Cortical magnification theory fails to predict visual recognition. European Journal of Neuroscience, 6, 1583–1588.

Strasburger, H., Rentschler, I., & Jüttner, M. (2011). Peripheral vision and pattern recognition: a review. Journal of Vision, 11(5), 1–82.

Strasburger, H., & Wade, N. J. (2015). James Jurin (1684–1750): A pioneer of crowding research? Journal of Vision, 15(1:9), 1–7.

Tolhurst, D. J., & Ling, L. (1988). Magnification factors and the organization of the human striate cortex. Human Neurobiology, 6, 247–254.

Tripathy, S. P., & Cavanagh, P. (2002). The extent of crowding in peripheral vision does not scale with target size. Vision Research, 42, 2357–2369.

Van Essen, D. C., Newsome, W. T., & Maunsell, J. H. R. (1984). The visual field representation in striate cortex of the macaque monkey: Asymmetries, anisotropies, and individual variability. Vision Research, 24(5), 429–448.

Virsu, V., & Hari, R. (1996). Cortical magnification, scale invariance and visual ecology. Vision Research, 36(18), 2971–2977.

Virsu, V., Näsänen, R., & Osmoviita, K. (1987). Cortical magnification and peripheral vision. Journal of the Optical Society of America A, 4, 1568–1578.

Virsu, V., & Rovamo, J. (1979). Visual resolution, contrast sensitivity and the cortical magnification factor. Experimental Brain Research, 37, 475–494.

Watson, A. B. (1987). Estimation of local spatial scale. Journal of the Optical Society of America A, 4(8), 1579–1582.

Wertheim, T. (1894). Über die indirekte Sehschärfe. Zeitschrift für Psycholologie & Physiologie der Sinnesorgane, 7, 172–187.

Westheimer, G. (1982). The spatial grain of the perifoveal visual field. Vision Research, 22, 157–162.

Whitney, D., & Levi, D. M. (2011). Visual crowding: a fundamental limit on conscious perception and object recognition. Trends in Cognitive Sciences, 15(4), 160–168.

